# Sex-biased gene expression precedes sexual dimorphism in the agonadal annelid *Platynereis dumerilii*

**DOI:** 10.1101/2024.06.12.598746

**Authors:** Rannyele P. Ribeiro, Ryan W. Null, B. Duygu Özpolat

## Abstract

Gametogenesis is the process by which germ cells differentiate into mature sperm and oocytes, cells essential for sexual reproduction. The sex-specific molecular programs that drive spermatogenesis and oogenesis can also serve as sex identification markers. *Platynereis dumerilii* is a research organism that has been studied in many areas of developmental biology. However investigations often disregard sex, as *P. dumerilii* juveniles lack sexual dimorphism. The molecular mechanisms of gametogenesis in the segmented worm *P. dumerilii* are also largely unknown. In this study, we used RNA sequencing to investigate the transcriptomic profiles of gametogenesis in *P. dumerilii* juveniles. Our analysis revealed that sex-biased gene expression becomes increasingly pronounced during the advanced developmental stages, particularly during the meiotic phases of gametogenesis. We identified conserved genes associated with spermatogenesis, such as *dmrt1*, and a novel gene *psmt*, that is associated with oogenesis. Additionally, putative long non-coding RNAs were upregulated in both male and female gametogenic programs. This study provides a foundational resource for germ cell research in *P. dumerilii,* markers for sex identification, and offers comparative data to enhance our understanding of the evolution of gametogenesis mechanisms across species.

**Summary statement:** This study provides insights into the mechanisms of gametogenesis in *Platynereis dumerilii* through comparative transcriptomics, unveiling sex-biased genes, including conserved and novel genes, governing this largely unexplored process.

## Introduction

Gametogenesis, the process of oocyte and sperm formation, is one of the fundamental processes of sexual reproduction, conferring genetic variability over generations and allowing species to propagate (Kikuchi and Tanaka, 2022; Nieuwkoop and Sutasurya, 1979; Padilla-Gamiño et al., 2022). This process involves a series of molecular events that orchestrate the differentiation of germ cells into mature gametes, either sperm or oocytes (Kikuchi and Tanaka, 2022). In gonochoric species, each individual produces only one type of gamete, while in hermaphroditic species, both sperm and eggs can develop in a single individual (Benvenuto and Weeks, 2020; Clark, 1978). Gonochorism is therefore characterized by distinct transcriptional programs that govern the development of sperm in males and oocytes in females, reflecting the divergent molecular requirements to determine and differentiate sex at the individual level (Kobayashi et al., 2012; Nagahama et al., 2021; Piferrer, 2021; Siebert and Juliano, 2017).

The segmented worm *Platynereis dumerilii* is a gonochoric species with unknown mechanisms of sex determination. The karyotype lacks heterochromosomes (Jha et al., 1995) that determine sex, such as in X/Y or Z/W sex determination systems (Smith, 2010; Wallis et al., 2008). In addition, no environmental cues have been identified thus far that determine sex in *P. dumerilii*, and laboratory cultures produce a male to female ratio approximated to 1:1 (Fig. S1A, B). Throughout most of their lifespan, these worms remain juvenile with no externally apparent morphologies that show sexual dimorphism until reaching maturity. As a part of sexual maturation, significant morphological changes occur, characterized by eye enlargement, loss of internal organs, and distinction of body regions: the atokous anterior body, with parapodia (segmental appendages) resembling those of juvenile stages, and the epitokous posterior body with specialized paddle-like chaetae (Fischer et al., 2010; Schulz et al., 1989; Tilic et al., 2023). Females, distinguished by their yellow coloration due to abundant eggs, have 20- 24 segments in the atokous body, while males exhibit a white atokous body with about 14-16 segments, and this difference is a part of sexual dimorphism exhibited by mature adults (Fig. S1C) (Schulz et al., 1989). On the other hand, juveniles exhibit no sexual dimorphism, appearing anatomically indistinguishable (Fig. S1D, E). While previous studies showed molecular differences between male and female juvenile brains (Schenk et al., 2019), to date there have been no molecular markers for identifying sex in individual juvenile worms without sacrificing the worms.

Gametogenesis in *P. dumerilii* has been predominantly characterized through histology. Both spermatogenesis and oogenesis take place within specialized tissues known as gonial clusters, which consist of groups of germ cells that are enveloped by a few or a single sheath cell (Fischer, 1974; Fischer, 1975; Meisel, 1990). Unlike conventional gonads, these gonial clusters are not confined to a specific location or organ but rather freely disperse within the coelom (the fluid filled internal cavity) of *P. dumerilii* (Fischer, 1974; Fischer, 1975; Meisel, 1990; Rebscher et al., 2007). Gonial clusters begin to populate the coelom in the trunk in juveniles, typically initiating in individuals with 35-40 segments (Kuehn et al., 2022).

This unique reproductive morphology of the worms where germ cells are spread across the body enables collection of the germ cells from worms at different developmental stages via amputating the posterior body. Because *P. dumerilii* can regenerate its posterior body after bisection, and because the bisection-regeneration process does not affect fertility (Metzger and Özpolat, 2024), these individuals then can be raised to sexual maturation, to determine whether they were males or females.

Therefore, it is possible to collect germ cell-enriched samples from the coelom during the juvenile phase when there is no sexual dimorphism, while keeping these worms alive to mature when they are finally sexually dimorphic.

In this study, we leveraged the ability of regeneration while staying fertile, and collected gonial clusters from worms at different developmental stages to cover different phases of gametogenesis, from early mitotic germ cell proliferation to different stages of meiosis. We then isolated RNA from each sample for bulk RNA sequencing and transcriptomic profiling to identify molecular mechanisms underlying gametogenesis and sex-biased expression patterns. We validated our results via whole-mount *in situ* hybridization chain reaction, and reverse transcription polymerase chain reaction. We found that sex-biased gene expression is dependent on the developmental stage, and we identified conserved and novel sex-related genes. This study serves as a foundational resource for comparative developmental biology, offering insights into the mechanisms of gametogenesis across annelids and lophotrochozoans in general. The mechanisms governing the proliferation and differentiation of germ cells in this species hold significant importance, particularly in the context of manipulating cellular differentiation programs in germ cells.

## Results

### RNAseq reveals sex-biased gene expression

To identify early differences between male and female juveniles, we focused on their germ cells, which undergo differentiation into sperm in males or oocytes in females during gametogenesis. Gametogenesis in *P. dumerilii* can be followed during the juvenile phase, well before the morphological changes associated with maturation arise. We conducted transcriptomic profiling of germ cell tissues in male and female juveniles, anticipating sex-biased gene expression related to gametogenesis and sex-differentiation (Fig. 1A).

**Fig 1.**
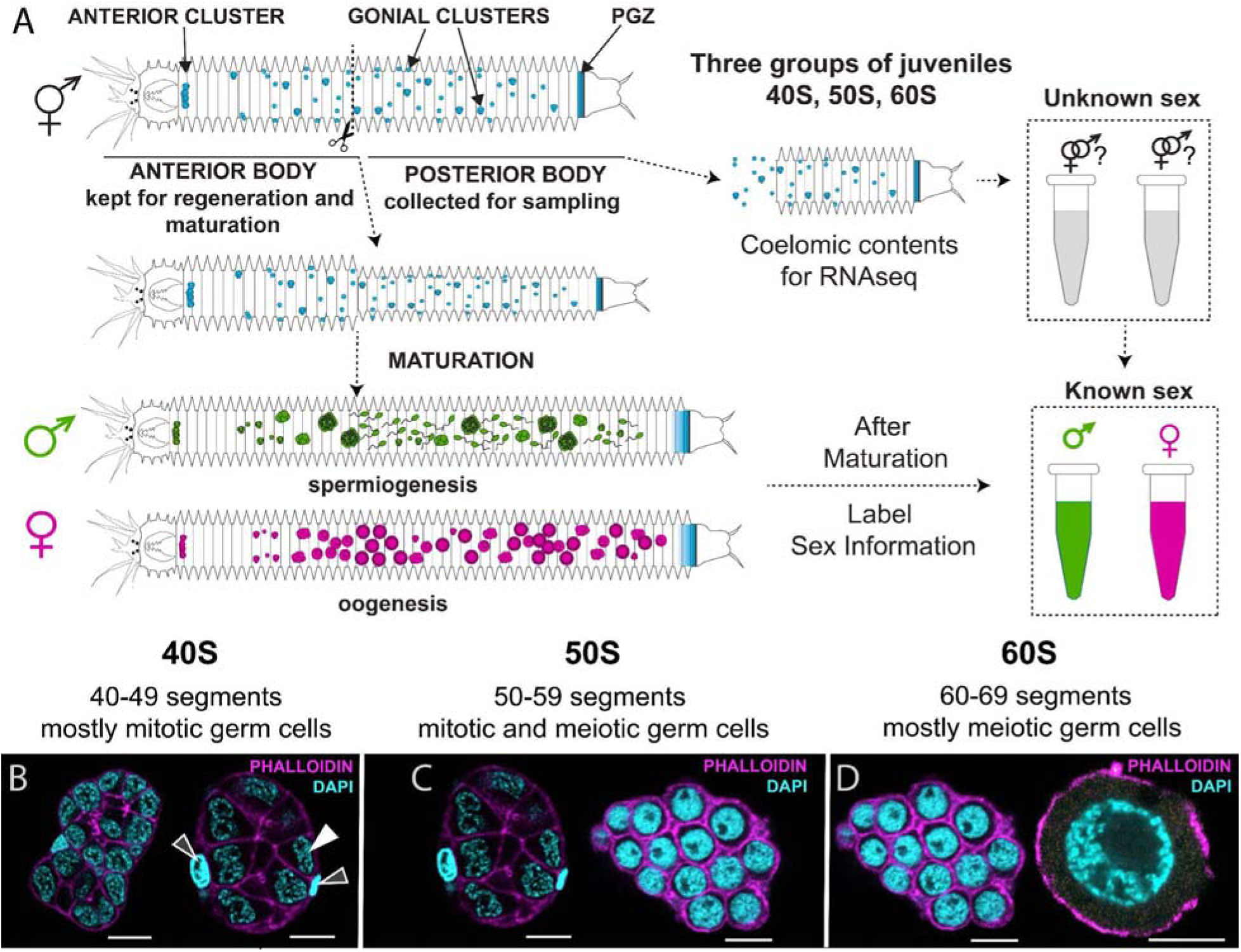
Experimental design for transcriptomic profiling of differentiating germ cells. (A) Gonial clusters, the cell aggregation that contains germ cells (shown in blue), were collected from the tails for RNA sequencing. Because juveniles do not show sexual dimorphism, the collected samples could not be sexed. Keeping the fragment with the head to regenerate the posterior body and further mature as male or female allowed us to obtain the sex information for each sample upon maturation of these individuals. The entire experiment lasted up to 10 months, until all worms matured. (B-D) We selected juveniles at different developmental stages, forming three groups: 40S, 50S and 60S. (B) 40S worms have 40-49 segments in length, and gonial clusters in mitotic stages mostly. The gonial clusters at this stage possess germ cells enveloped for 1-3 sheath cells. Solid white arrowhead points to a sheath cell, outlined white arrowheads point to germ cells. (C) 50S worms have 50-59 segments and their gonial clusters are abundant, including both at mitotic and meiotic stages of gametogenesis. (D) 60S worms have 60-69 segments and clusters in meiotic stages mostly (see Fig. 6D for details), but mitotic gonial clusters may be present. Scale bars: 10µm.

The germ cells of *P. dumerilii* are organized into clusters surrounded by sheath cells. We refer to these structures as gonial clusters (Metzger and Özpolat, 2024; Rebscher et al., 2007; Rebscher et al., 2012). These gonial clusters do not have a specific localization but rather spread across the body in the coelomic cavity.

Transitions in gonial cluster stages are observed over the course of juvenile growth. The worms grow by segment addition from a posterior growth zone (PGZ) localized at their tails throughout their lifespan (Balavoine, 2014; Gazave et al., 2013). Once the worms surpass about 35 segments in growth, these gonial clusters can be collected from the trunk (Kuehn et al., 2022). The number of gonial clusters in the trunk increase, as the worms grow by adding more segments and eventually mature worms with differentiated gametes being typically 70-80 segments long.

The gonial clusters are present in the coelom alongside other cell types, such as coelomocytes. Because of the challenges in isolating gonial clusters and the potential involvement of surrounding cells in gametogenesis, we collected coelomic contents for RNA sequencing, therefore, our datasets are enriched for gonial clusters, but also include some other coelomic cell types. Our strategy involved a bisection-regeneration approach which allowed us to collect gonial clusters from samples while keeping the head segments alive so we could sex them once they regenerated and matured (Fig. 1A). We selected three groups of worms that represent the transitions of gametogenesis and developmental stages, based on gonial cluster staging (Fischer, 1974; Fischer, 1975; Meisel, 1990) and worm size measured by segment number (Kuehn et al., 2022). These groups were: **40S**, worms with 40-49 segments that have mostly mitotic germ cells in gonial clusters (Fig. 1B); **50S**, worms with 50-59 segments that have a combination of mitotic and early meiotic germ cells (Fig. 1C); and **60S**, worms with 60-69 segments that have germ cells more progressed in gametogenesis, mostly at meiotic stages (Fig. 1D, Supplementary Material 1). Next, we bisected the worms and used the posterior bodies to sample coelomic contents for bulk RNAseq (Fig. 1A). Each sample yielded approximately 50-100µL of coelomic contents, including several gonial clusters and coelomocytes in buffered medium (Nereis Balanced Saline Solution, NBSS). A total of 30 coelomic content samples were collected and sequenced.

To generate a reference transcriptome for further analysis, we selected three samples for each sex (male and female) and group (40S, 50S, 60S), a total of 18 samples (Supplementary Material 1). The reference transcriptome contained 727,308 transcripts, covering 300,594 genes (“Trinity” genes) with a high completeness (96.5%) (Table 1). This reference transcriptome was used for transcript quantification and differential expression analysis. We performed an in-depth differential expression (DE) analysis at the gene level, using a total of 30 samples (5 replicates per each sex and group). This analysis revealed a total of 424 genes with differential expression, with 122 genes up-regulated in males and 302 genes up-regulated in females (p <0.001) (Fig. 2A).

**Fig. 2.**
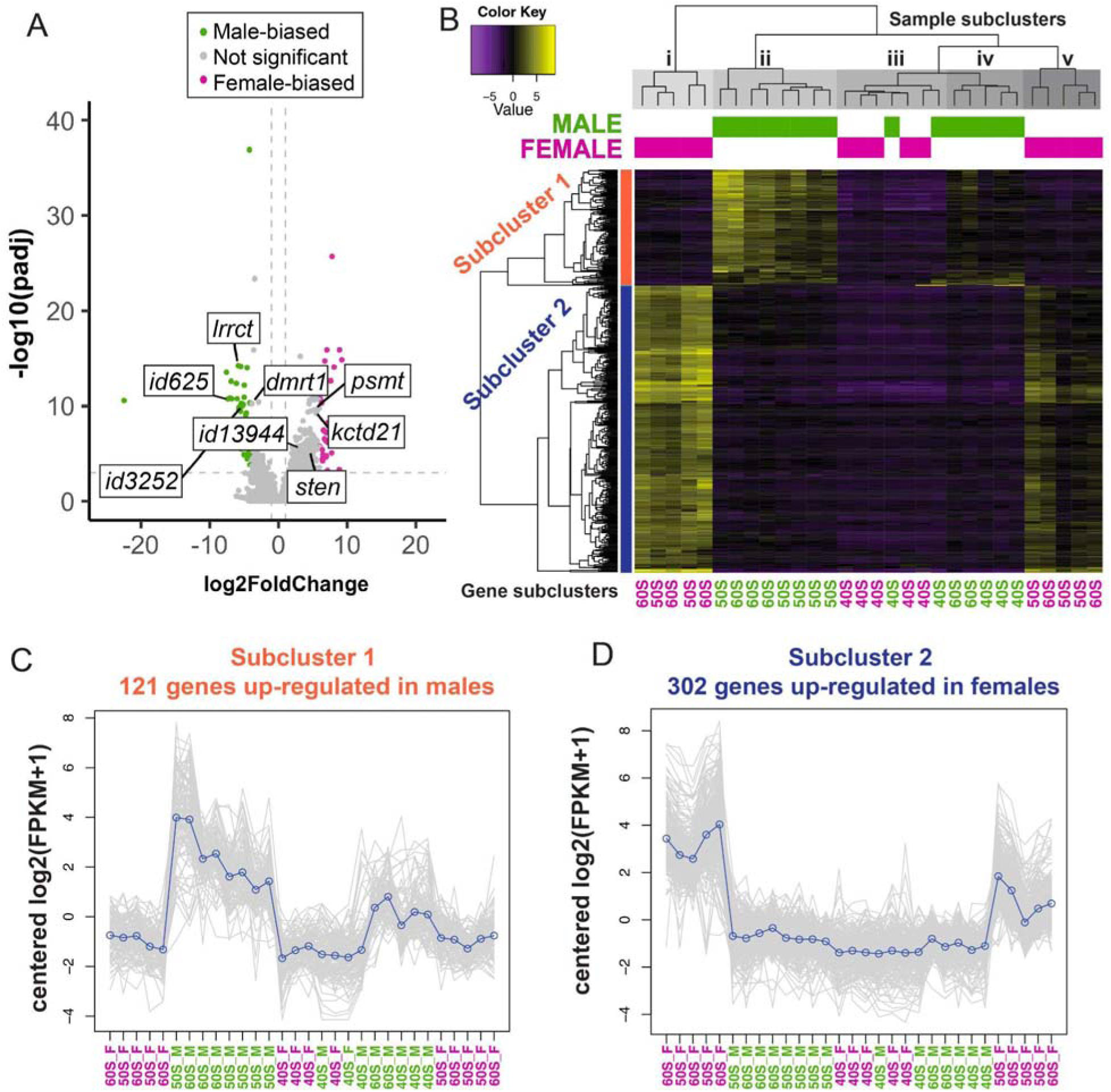
Sex-biased gene expression in *P. dumerilii* juveniles. Results of the differential gene expression analysis at the level of Trinity genes. (A) Volcano plot shows the distribution of Trinity “genes” according to their log-to-fold change (log2FoldChange). The analysis used “female” as a reference condition, therefore, negative values represent genes up-regulated in males, while positive values represent genes up-regulated in females. (B) Heatmap of differentially expressed genes over 30 samples, including 5 replicates for each sex and group (40S, 50S and 60S). Samples were clustered by similarity in gene expression patterns. There are two groups of females (subclusters I and V) and two groups of males (subclusters II and IV), while males and females of 40S, the younger group of worms (mostly females), cluster together (subcluster III). (C-D) Subclusters 1 and 2 of differentially expressed (DE) genes. (C) Subcluster 1 shows genes up-regulated in males. (D) Subcluster 2 includes all DE genes up-regulated in females. A third subcluster containing one gene up-regulated in males was generated by the analysis and is available in the Fig. S2.

**Table 1.** Statistical summary raw data, transcriptome assembly, and functional annotation of *P. dumerilii*’s germ cell transcriptome.

| Parameters | Value |
| --- | --- |
| Raw reads* | 391.17 GB |
| Number of clean reads | 345.56 GB |
| Total assembled bases | 524,872,716 |
| Median contig length (nucleotides) | 387 |
| Average contig length (nucleotides) | 721.66 |
| N50 value (nucleotides) | 1160 |
| Total number of transcripts | 727,308 |
| Percentage of GC | 40.75 |
| Trinity 'genes' | 300,594 |
| Predicted proteins | 158,401 |
| Transcripts with GO annotation | 25,269 |
| Completeness | 96.5% |
| Duplicated copies | 87.9% |
| Single copies | 8.6% |
| Fragmented copies | 10 |

\*Sum of raw reads used for the reference transcriptome assembly.

Next, we compared the expression patterns of DE genes by generating a heatmap. This visualization approach allowed us to analyze the expression profiles of these DE genes across the different samples, revealing similarities and differences. The heatmap clearly highlighted a sex-biased gene expression, particularly in the 50S and 60S groups (Fig. 2B). It revealed two main clusters of female samples (i and v) and two clusters of male samples (ii and iv), primarily separating the 50S and 60S worms (Fig. 2B). Another subcluster (iii) included the 40S male and female samples, showing a general downregulation (mostly negative values), and clustering with the male sample subcluster iv (Fig. 2B). The analysis also generated subclusters of DE genes, which were grouped into two main subclusters: one containing 121 DE genes were up- regulated in males, subcluster 1 (Fig. 2B, C, Fig. S2A-E), and another with 302 genes up-regulated in females, subcluster 2 (Fig. 2B, D, Fig. S2A-E).

Our differential gene expression analysis significantly indicates overall sex-biased transcription profiles in juveniles of *P. dumerilii*, with more pronounced differences in 50S and 60S worms. These developmental stages represent the transitions from mitotic to meiotic stages of gametogenesis, as evidenced by previous research (Fischer, 1974; Fischer, 1975; Meisel, 1990).

### GO enrichment analysis indicates gametogenesis-related genetic pathways

To determine the underlying biological mechanisms and functions associated with the identified DE genes, we conducted a Gene Ontology (GO) enrichment analysis. This analysis categorized gene functions into three main domains: biological processes, molecular functions, and cellular components, along with their respective subcategories. We integrated the transcriptome annotation report to the analysis of differential expression between males and females, and highlighted the top 20 enriched GO subcategories (Fig. 3A, B). The GO enrichment analysis revealed that the DE genes up- regulated in males were significantly enriched for subcategories related to spermatogenesis and sperm function such as "sperm principal piece", "sperm motility", "sperm capacitation" and "CatSper complex" (Fig. 3A). DE genes up-regulated in females were enriched for GO subcategories associated with metabolic processes, for example, "primary metabolic process", "macromolecule metabolic process" and "organic substance metabolic process" (Fig. 3B). Additionally, categories of the domain cellular component were enriched in females, including "organelle", "intracellular membrane" and "non-membrane organelle" (Fig. 3B). The GO enrichment analysis supports that our transcriptomes accurately represent profiles expected for germ cells in gametogenesis, and indicate sex-biased differences.

**Fig. 3.**
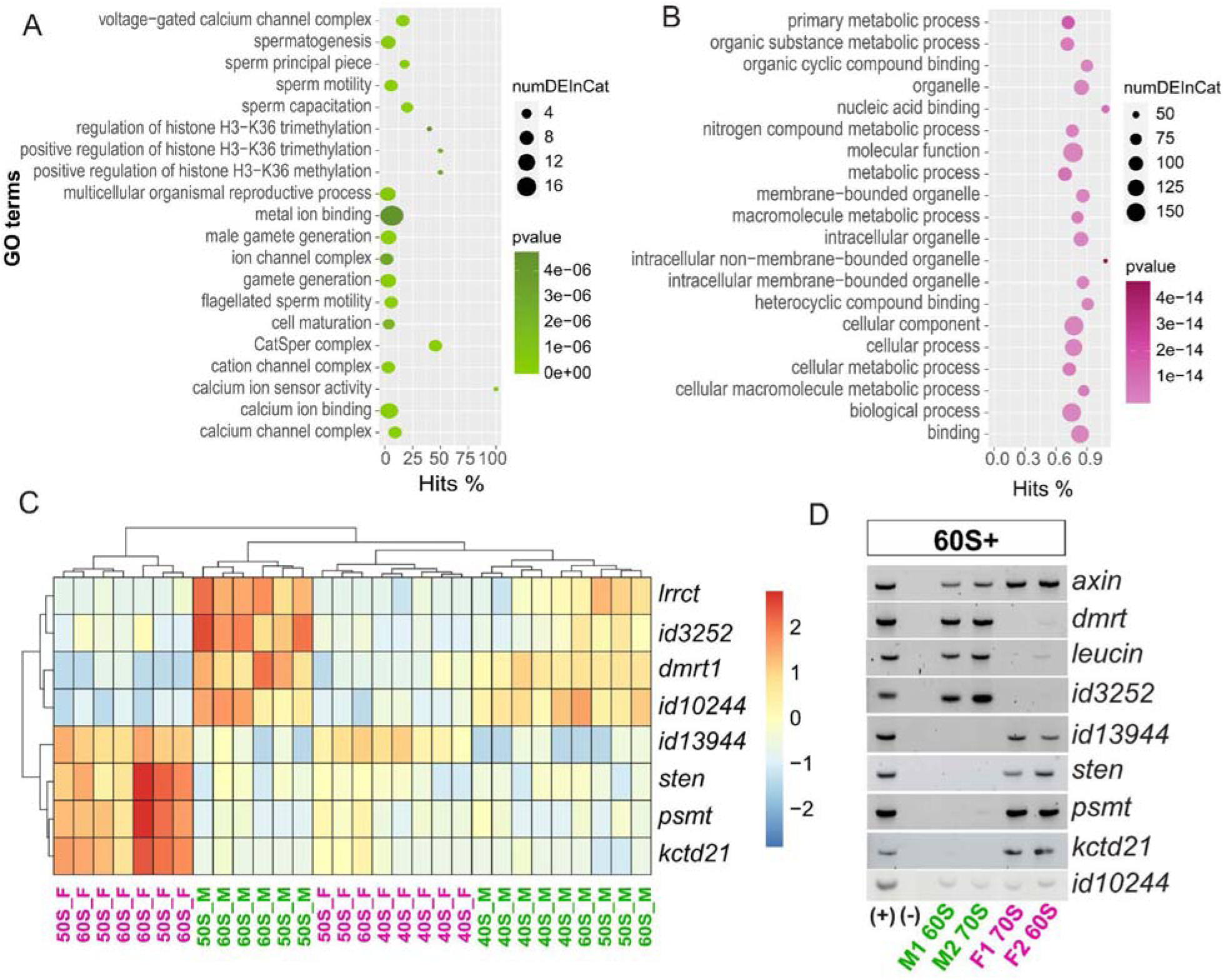
Gene ontology (GO) enrichment and validation of selected differentially expressed (DE) genes. (A-B) Top 20 enriched GO categories in DE genes up-regulated in males (A) and females (B), ranked from top to bottom. (C) Expression patterns of DE genes selected for validation in a heatmap across 30 samples of 40S, 50S and 60S males and females. (D) RT-PCR results of the validated DE genes in agarose gels for samples of two females and two males. 60S, worms with 60-69 segments. 70S, worms with 70-79 segments. Additional RT-PCR results and quantification of gene expression are provided in Supplementary Figure S3.

Finally, our experimental design made the assumption that there is no sex change resulting from bisection and regeneration. In our previous experiments we have not observed that sex changes after regeneration in *P. dumerilii* (Metzger and Özpolat, 2024). To ensure that sex indeed does not change after regeneration, we bisected 37 worms with 40-50 segments and cultured them individually. We determined their sex before and after regeneration based on the gonial cluster morphology (Fischer, 1974; Fischer, 1975; Meisel, 1990) as well as using molecular male and female markers determined via our RNAseq experiment (see below, and Methods). Based on these analyses, we found that none of these worms exhibited sex change after regeneration (Supplementary Material 2).

### Validation of sex-biased gene expression

To verify the biological reproducibility of the observed transcriptomic differences, we performed reverse transcription polymerase chain reaction (RT-PCR) and *in situ* hybridization chain reactions (HCRs) using gonial clusters or whole worms, respectively. RNAseq may present “noisy” transcripts resulting from technical issues (Varabyou et al., 2021); therefore, validating selected genes can provide a more accurate picture of their gene expression patterns. Additionally, if the selected genes can show differences using quick techniques like RT-PCRs, they can be good sex markers that can be incorporated to any experimental setup for *P. dumerilii*. We selected 12 DE genes among those ranking at the top 50 differentially expressed genes as for the adjusted p-value (0.001) (Supplementary Material 3). These genes were selected based on a cross reference comparison between two differential expression analysis: a preliminary analysis with 18 samples at isoform level, and an in-depth analysis with 30 samples at gene level (Supplementary Materials 4-6, see Methods for more details).

We conducted RT-PCRs on coleomic samples using the same collection method as for our RNAseq, to test the expression patterns of the selected DE genes. PCR primers were designed using the transcript isoforms. RT-PCR was successful for 8 of the 12 target genes (Supplementary Material 7). These included four genes upregulated in males: *dmrt1* (double-sex mab-3 transcription factor 1), *lrrct* (leucine rich repeat C terminal), *id3252*, *id10244*; and four genes upregulated in females: *sten* (Secretory 24CD), *kctd21* (Potassium channel tetramerization domain containing 21), *psmt* and *id13944*. A heatmap generated using data from the 30 samples in the in-depth analysis showed the expected sex-biased expression of these 8 genes in juvenile worms (Fig. 3C).

To validate these results, we conducted RT-PCRs in samples from worms with more than 60 segments, which were sexed based on gonial cluster morphology. The RT-PCR assays exhibited the expected expression patterns in the 60S worms for most genes, with three male up-regulated genes (*dmrt1*, *lrrct*, *id3252*) showing higher expression in male samples, and the four female up-regulated genes (*id13944*, *sten*, *psmt*, *kctd21*) showing higher expression in female samples (Fig. 3D, Fig. S3).

However, *id10244*, which was up-regulated in males from the RNAseq, did not show sex-biased expression in the RT-PCRs (Fig. 3D).

We also targeted 40S worms for RT-PCRs, from which we collected samples of coelomic contents from the tails, keeping the anterior body to regenerate and grow.

These samples were used to examine the expression of three differentially expressed (DE) genes identified from our RNAseq data: *dmrt1*, which was up-regulated in males, *psmt* and *id13944*, both of which were up-regulated in females. The overall success rate of the RT-PCRs was much lower in 40S worms, with only 4 out of 20 reactions successfully amplifying the housekeeping gene (Fig. S3C, Supplementary Material 7).

We investigated two of these worms, labeled as A40 (male) and A41 (female). A40 showed higher expression of *dmrt1* compared to *psmt* and *id13944* as a 40S worm based on the RT-PCRs (Fig. S3C), while A41 showed higher expression of *psmt* and *id13944* compared with *dmrt1* (Fig. S3C). As such worms were kept to regenerate and grow, when they reached the stage of 60S, we used the same worms for *in situ* HCRs to visualize their gonial clusters and corresponding gene expression for *dmrt1* and *psmt* (Fig. 4A, B). Consistent with RT-PCRs, the male sample showed expression of *dmrt1*, and no expression of *psmt* at the 60S stage (Fig. 4A-D, A’-D’). Interestingly, the female sample exhibited expression of not only the expected *psmt* but also *dmrt1* at the 60S stage (Fig. 4E-H, E’-H’).

**Fig. 4.**
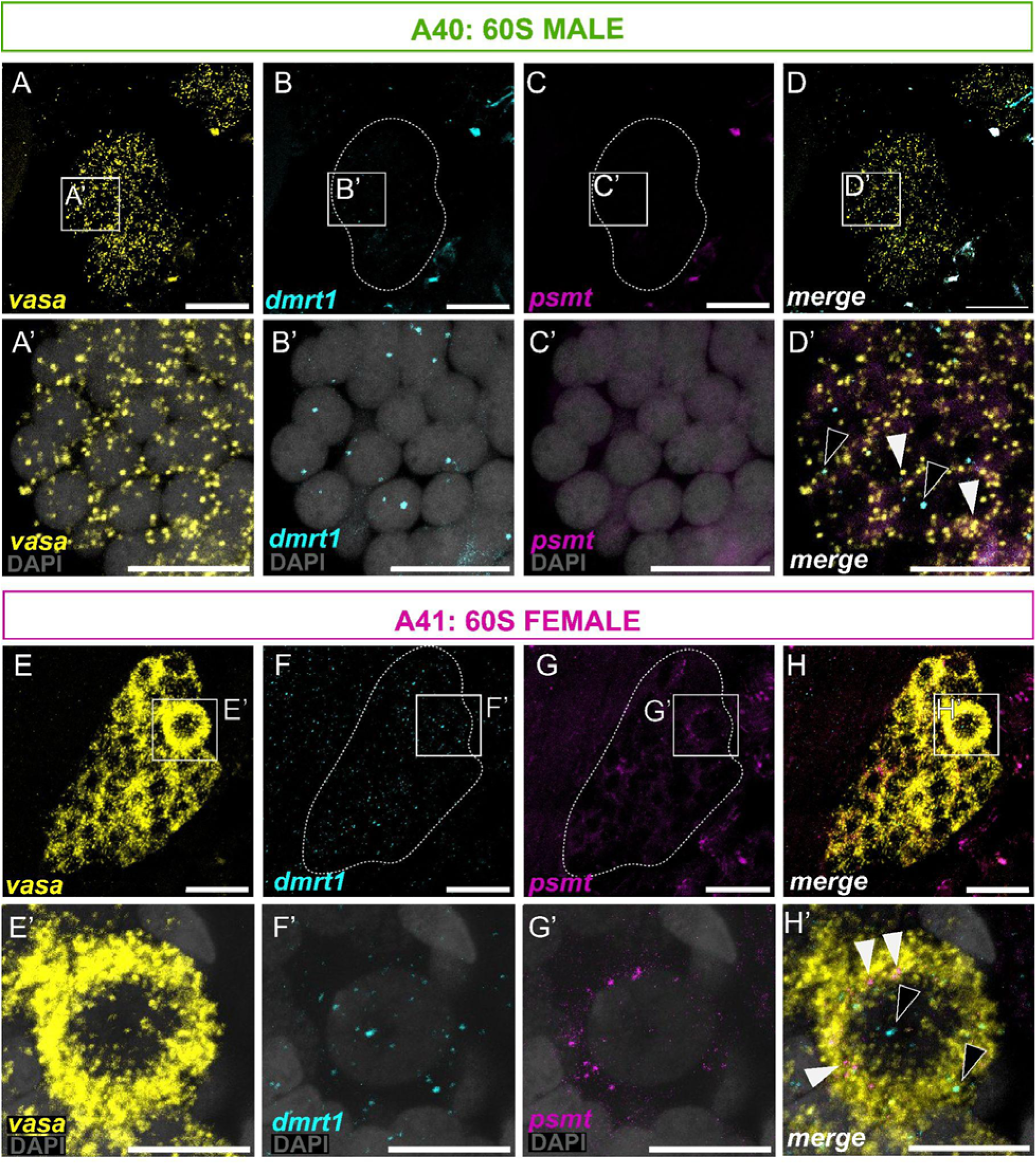
Expression of *dmrt1* and *psmt* in gonial clusters. (A-D, A’-D’) Gonial cluster from a male 60S worm. This gonial cluster is in stage 3, spermatocyte ball and clearly shows expression of *dmrt1* (DE gene up-regulated in males, B, B’) and absence of *psmt* (DE gene up-regulated in females, C, C’). (E-H, E’-H’) Gonial cluster from a female 60S worm. Its morphology is primarily of stage 2 (synaptic oocytes) but also exhibits some late prophase I oocytes with larger nuclei (stage 3). Expression of both *dmrt* and *psmt* is noted (F, G, F’, G’). (E’-H’) Close-up of a late prophase I stage 3 oocyte showing expression of both *dmrt1* and *psmt*. Dashed lines mark the borders of the gonial clusters. These gonial clusters are from worms initially assessed via RT-PCRs when at 40S (Fig. S3). Asterisks indicate background (Fig. S4). Scale bars: A-H 20µm, A’-H, 10µm.’

Overall, as a detection method, RT-PCRs seemed to be more reliable for 60S worms, while 40S worms presented technical challenges. The expected sex-biased gene expression evidenced by our RNAseq was largely validated by RT-PCRs, except for the gene *id10244*. Interestingly, while *dmrt1* expression was detected only in males by RT-PCRs, HCRs revealed *dmrt1* expression in both males and females, possibly due to differences in expression levels in either sex.

### Sex-differentiation genes are expressed in gonial clusters

While RT-PCRs were instrumental in validating the DE genes in samples of coelomic contents, they did not provide spatial information. In addition, given that the coelomic contents include both gonial clusters and coelomocytes, RT-PCRs cannot confirm sex-biased expression in gonial clusters alone. To address this, we used *in situ* hybridization chain reaction (HCR) to verify whether DE genes express in gonial clusters. We performed HCRs in whole worms ranging from 30 to 70 segments, using *vasa* expression as a positive marker for gonial clusters (Kuehn et al., 2022; Metzger and Özpolat, 2024; Rebscher et al., 2007; Zelada Gonzáles, 2005). We tested HCR oligo pools for the selected 12 DE genes from our RNAseq data (Supplementary Material 3). HCRs were successful for a total of 8 genes, 4 of the DE genes up-regulated in males (*dmrt1*, *lrrct*, *id635*, and *id3252*) and 4 of the DE genes up-regulated in females (*id13944*, *sten*, *psmt*, *kctd21*).

We observed that all these DE genes can be expressed in both male and female gonial clusters (Fig. 5A, B, Supplementary Material 8). However, we noticed a tendency for the male-upregulated genes to be enriched in male gonial clusters (*dmrt1*, *lrrct*, *id3252*, *id625*) (Fig. 5B, Fig. S5), while the female-upregulated genes were more frequent in female gonial clusters (id13944, *sten*, *psmt*, *kctd21*) (Fig. 5B, Fig. S6). To quantify our *in situ* HCR results, we counted the presence and absence of each gene in several gonial clusters in each sample (see Statistics in Methods). The data on the presence and absence was used to calculate the odds ratio of male-to-female expression. The odds ratio provided a measure of how likely (or unlikely) it is to observe the expression of a given gene in male gonial clusters compared to the female ones.

**Fig. 5.**
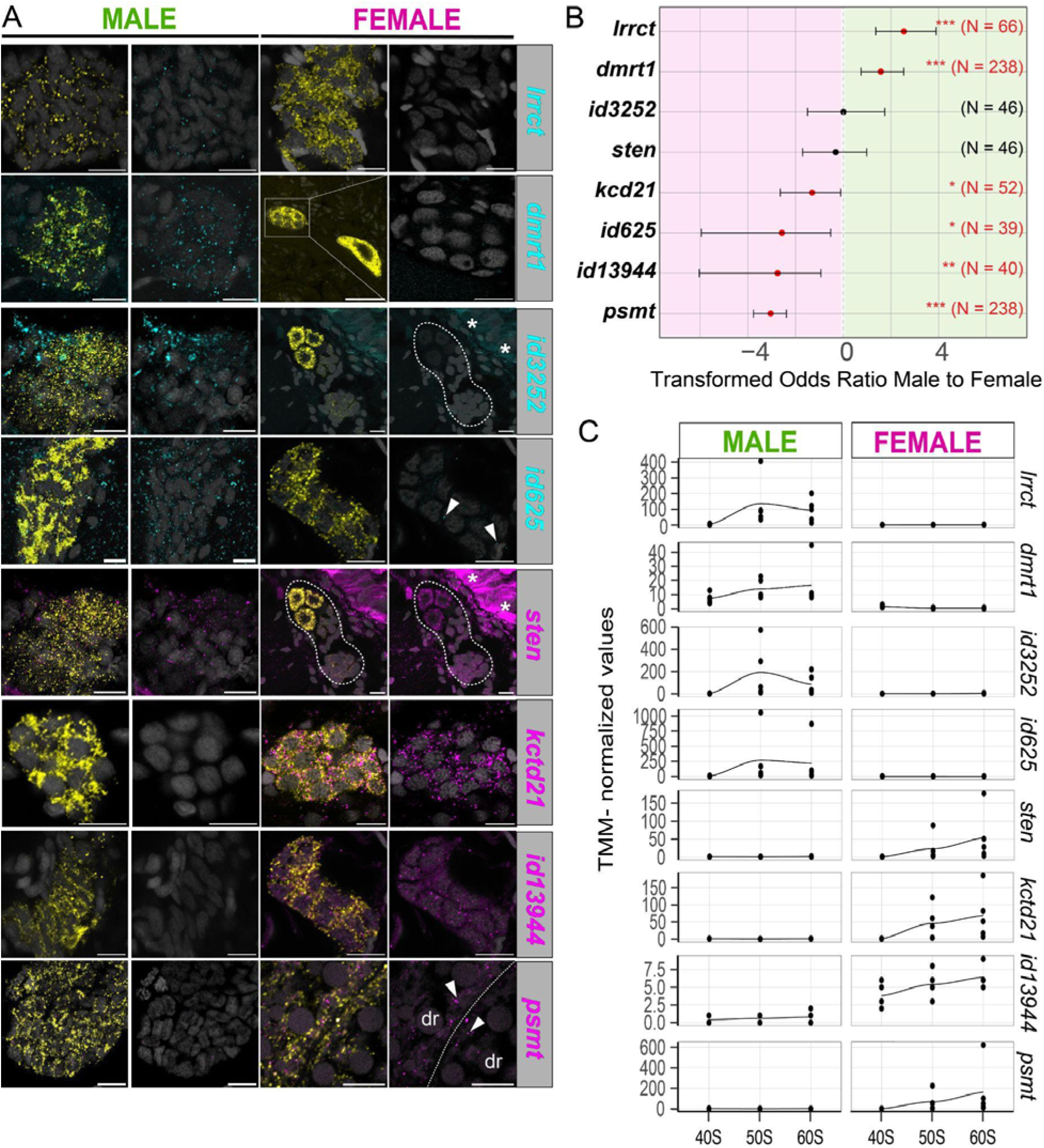
Gene expression patterns of DE genes in gonial clusters. (A) Whole-mount *in situ* hybridization chain reaction performed on both male and female gonial clusters (50S and 60S worms). *vasa* expression is shown in yellow, male-biased genes in cyan, and female-biased genes in violet. RNA-seq revealed *lrrct*, *dmrt1*, *id3252*, and *id625* as male-biased genes. In situ hybridization images show their expression in male gonial clusters and absence in female gonial clusters. For *id625*, only a few expression sites are observed in the female gonial cluster, marked by white arrowheads. The analysis identified *sten*, *kcdt21*, *psmt*, and *id13944* as female-biased genes. Images demonstrate their expression in female gonial clusters and absence in male gonial clusters, except for *sten* which also expresses in male gonial clusters. In females, *psmt* is expressed in vitellogenic oocytes near the cellular membrane, with arrowheads indicating the expression sites. Dashed lines demarcate the borders between two oocytes, and droplets are noted (dr). Detailed expression patterns of *dmrt1* and *psmt* can be found in Figs. S5 and S6. Asterisks denote background signals. Scale bars = 10µm, except female panels for *dmrt1* and *psmt*, which are 20µm. (B) The male-to-female odds ratio of gene expression in gonial clusters, with N representing the number of gonial clusters analyzed. Positive values indicate a higher probability of detecting gene expression in male gonial clusters (green right side), while negative values indicate a lower probability (vice-versa for females, pink left side) (p<0.05). Hence, positive tendency characterizes a male-biased gene (green zone), while negative tendency characterizes a female-biased gene (pink zone). The expression of *id625* tends to be present in a large number of gonial clusters, showing a female-biased pattern. However, the expression of *id625* is usually low, as observed in HCRs (A). Number of males (Nm) and females (Nf): *lrrct*, Nm=6, Nf=7; *dmrt1*, Nm=15, Nf=15; *id3252*, Nm=7, Nf=2; *sten* Nm=7, Nf=2; *kcd21*, Nm=7, Nf=4 ; *id13944*, Nm=4, Nf=6; *psmt*, Nm=6, Nf=7. (C) Transcript quantification of selected DE genes displayed in panel A.

This analysis revealed that at least two (*dmrt1* and *lrrct*) out of four male-upregulated genes were more likely to be expressed in male gonial clusters, based on the 95% confidence intervals (Fig. 5B). Out of four female-upregulated genes, three genes (*id13944*, *psmt*, and *kctd21*) showed odds ratios favoring female gonial clusters (Fig. 5B). Interestingly, *id625*, a gene found to be upregulated in males from the RNAseq, appeared more likely to be expressed in female gonial clusters. *id625* and *id3252* showed no clear sex-bias in gonial cluster expression based on the odds ratio analyses of our HCR results (Fig. 5B).

The RNAseq data provided a global assessment of differential gene expression, which does not necessarily mean that DE genes were exclusively expressed in one sex or another. Upon further examination of the RNAseq read counts, we found that while there were substantial differences in expression between males and females in TMM normalized values of expression (Fig. 5C), these genes have detectable levels of expression in both sexes using HCRs (Fig. 5A).

The HCRs revealed sex-biased expression for a subset of the DE genes within the gonial clusters. We consider *dmrt1* and *lrrct* to be reliable male-related genes, while *kctd21*, *id13944*, and *psmt* appear as reliable female-related genes. However, the sex-biased expression of *id3252*, *sten*, and *id625* remained inconclusive as robust sex markers due to the discrepancies between the RNAseq, RT-PCR, and HCR results.

### Anterior cluster and posterior growth zone do not show sex-biased gene expression

In addition to gonial cluster expression, we also analyzed expression of the sex-biased markers in other structures and tissues in our samples. Since we had *vasa* as a germline/multipotency program marker (Gazave et al., 2013; Kuehn et al., 2022; Rebscher et al., 2007; Zelada Gonzáles, 2005), we were able to assess the expression of the DE genes not only in the gonial clusters, but also in other *vasa*-positive tissues, such as the anterior cluster (AC), a group of *vasa*-positive cells near the head, and the posterior growth zone (PGZ), located near the tail. We found that most of the DE genes selected are expressed in both AC and PGZ (Fig. 6A-C) with no sex-biased expression, as evidenced by the male-to-female odds ratios (Fig. 6B-C, Fig. S7). Therefore, the sex-biased expression observed for the DE genes appears to be specific to the gonial clusters only and not present in other *vasa*-positive tissues.

**Fig. 6.**
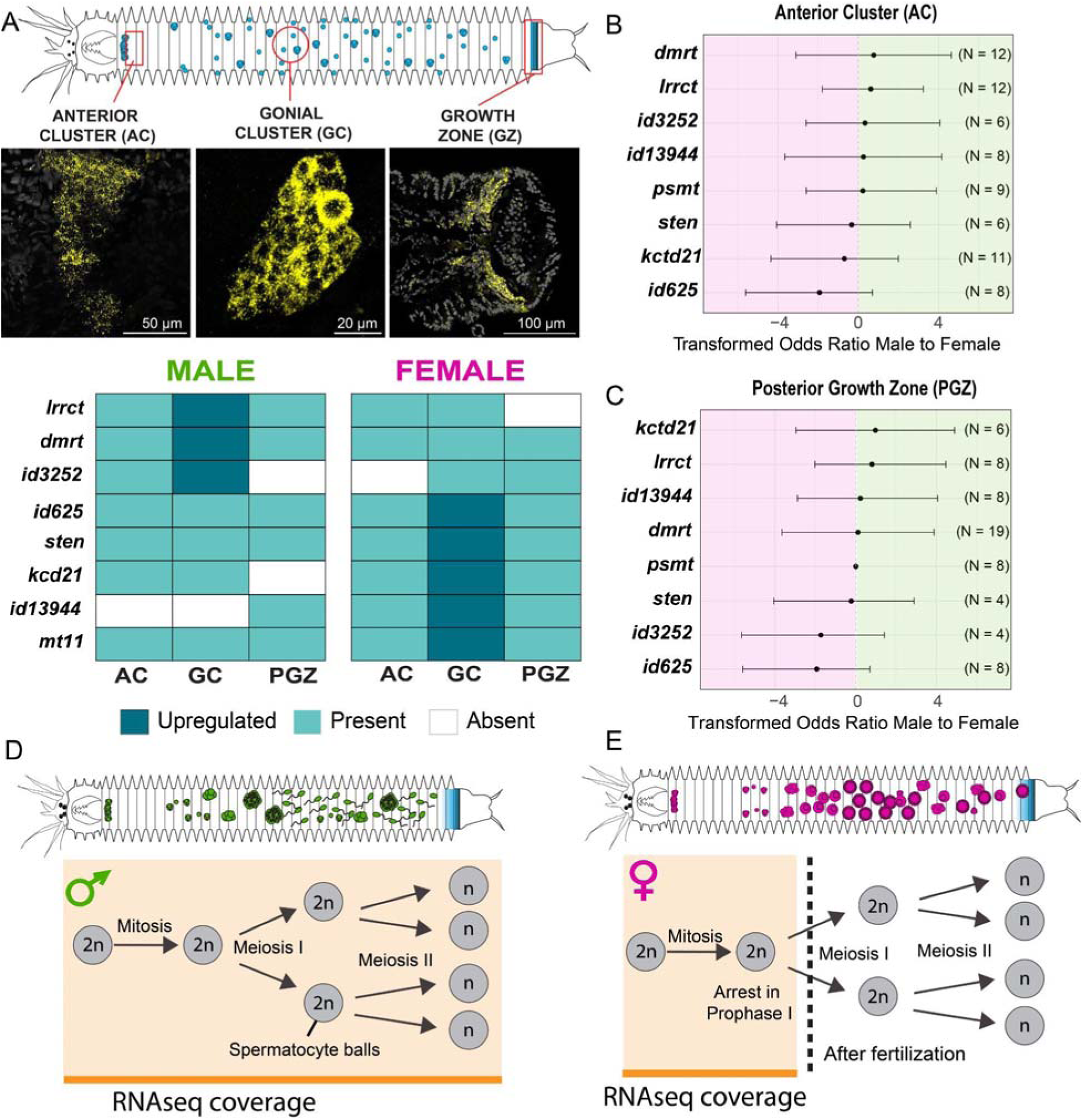
Expression patterns of sex-related genes in germline multipotent-like tissues. (A) Summary of expression patterns of selected DE genes via HCRs. Gonial clusters show quantitative sex-biased differences, while anterior cluster and posterior growth zone show no significant sex-biased differences. (B-C) Male-to-female odds ratio for anterior cluster (p<0.05) (B) and posterior growth zone (C), positive tendency characterizes a male-biased pattern (green right zone), while negative tendency characterizes a female-biased pattern (pink left zone). (D) RNAseq coverage of the gametogenesis stages in males. We captured spermatogenesis up to stage 3 and 4, spermatocyte balls and sperm. (E) RNAseq coverage from female gonial clusters. Gametogenesis is arrested in prophase I early in gametogenesis and is rescued only after fertilization.

### In-depth analysis of dmrt1 and comparison with other animals

This study identified *dmrt1* upregulated in male gonial clusters, as evidenced by RNAseq, RT-PCRs, and HCR. *dmrt1* was annotated using the Trinotate pipeline and a comprehensive phylogenetic analysis. Our phylogenetic tree, based upon an alignment of the double-sex mab-3 (DM) domains, robustly places *dmrt1* transcript within the *dmrt1* clade, supported by Shimodaira and Hasegawa approximate Likelihood Ratio Test (SH-aLRT = 65.6), approximate Bayesian Likelihood (0.944), and Ultrafast bootstrapping statistics (79) (Fig. S8, S9). The DM domain of *P. dumerilii -* DMRT1 predicted protein was further compared to an AlphaFold2 prediction of the Human DMRT1 crystal structure (PDB: 4YJ0) (Fig. S10). Key cysteine and histidine residues that coordinate zinc ion binding are positionally conserved between the *P*. *dumerilii* and human structures when the two are superimposed (Fig. S10).

While we observed quantitative differences of *dmrt1* expression between males and females, it is plausible that certain *dmrt1* isoforms are preferentially expressed in females rather than in males (Clough et al., 2014). To investigate this, we captured and analyzed all isoforms of *dmrt1* from our RNA-seq data, and found that all isoforms are predominantly expressed and are up-regulated in males (Fig. S11), therefore we did not detect any female-specific isoforms.

In addition, because other genes with DM domain can be up-regulated in females, such as *dmrt5* in zebrafish ovaries (Graf et al., 2015), we looked into other *dmrt*-like genes in *P. dumerilii*. Besides *dmrt1, P. dumerilii* presents other three genes with DM domain so far, including *dmrt1/2*, *dmrt2*, and *dmrt3* according to our gene trees (Fig. S8). In our study only *dmrt1* has been detected as a DE gene, at the our selected level of significance (FDR<0.001).

DMRT1 protein is associated with members of the SOX genes, particularly SOX9 in other organisms. DMRT1 facilitates chromatin accessibility, enabling SOX9 binding regions associated with male sexual development in mammals (Lindeman et al., 2021). In our transcriptome, we identified four potential *sox* genes, and found that *sox2* was slightly up-regulated in females with a log2 fold change of 0.75, although it did not meet our differential expression significance threshold. Indeed, in RT-PCRs for *sox2* we found similar levels of expression in both males and females (Fig. S3, Fig. S12 for gene tree). HCRs also show expression of *sox2* in male and female gonial clusters, anterior cluster and posterior growth zone (data not shown). We also found *sry*, another member of the SOX family associated to male specification (Clough et al., 2014), upregulated in males, according to our preliminary differential analysis, ranking 47778 in our in-depth analysis, disqualifying it as a DE gene (Suppl. Materials 5; Fig. S12 for SOX tree). Therefore, both *sox2* and *sry* were removed from our selected list of DE genes.

### dmrt1 is a male-biased gene while sten indicates tissue-specific sex-biased expression patterns

Previous studies have found *dmrt1* and *sten* to be up-regulated in the brain of mature males in *P. dumerilii* based on RNAseq data (Schenk et al., 2019). To determine if this pattern holds true in juveniles, we analyzed *dmrt1* and *sten* expression patterns in the brain, using whole-mount HCR. Interestingly, we found no differences between juvenile male and female brains: both *dmrt1* and *sten* were expressed at low levels (Fig. S13). Therefore, this study highlights *dmrt1* as a male-biased gene for gonial clusters of *P. dumerilii*. Remarkably, while *sten* expression appears biased towards the male brain in mature worms (Schenk et al., 2019), our RNAseq and RT-PCR assays revealed prominent *sten* expression in gonial clusters of female juveniles. As mentioned above HCRs did not reveal significant sex-biased expression of *sten* within the brains of juveniles younger than the ones tested in previous studies, suggesting that the sex- sepecific expression differences in the brain arise closer to sexual maturation. Together, these results underscore the tissue-specific nature of sex-biased gene expression.

### psmt is a protostome-specific methyltransferase

Our RNAseq revealed *psmt* as a gene up-regulated in female gonial clusters. Initially it could not be annotated with confidence to a certain gene, but BLAST results pointed to a type of methyltransferase. To confirm the ontology of this gene as a methyltransferase, we generated a gene tree. The gene tree revealed that *psmt* is a methyltransferase specifically present in protostomes (Fig. S14) and clusters with a group of S-adenosyl-L-methionine (SAM) dependent methyltransferases, as supported by Shimodaira and Hasegawa approximate Likelihood Ratio Test (SH-aLRT = 99.6), approximate Bayesian Likelihood (1), and Ultrafast bootstrapping statistics (97) (Fig. S15). AlphaFold2 modeling showed a good fit to the overall folds of several SAM- dependent methyltransferase crystal structures (Fig. S16). This included the ligand binding pocket where both the methyl donor, SAM, and its post-donation product, S- adenosyl-L-homocysteine (SAH) were comparable to bona fide methyltransferases (Fig. S17). The 50 amino acids of the N-terminus had weak confidence (lDDT score < 60), but this same region also had only a few reference sequences to be modeled upon (Fig. S16). Further analysis flagged this region (from Cysteine-15 to Proline-37) as being a transmembrane domain. In summary, our analyses suggest that *psmt* encodes a SAM- dependent methyltransferase present across protostomes including *P. dumerilii*. Its female-biased expression in *P. dumerilii* gonial clusters hints at a potential role for this gene in female sexual development or oogenesis.

### Putative long non-coding RNAs show sex-biased expression

Our annotation pipeline has not identified the ontology of the transcripts *id3252* and *id13944*. These uncharacterized transcripts contain short open reading frames (ORFs), with <150 bp, and considerable length, with *id3252* having 400 bp and *id13944* having 834 bp. Initially, we speculated that they were artifacts of our assembly. However, this was ruled out following successful RT-PCR validation and sequencing of amplicons matching their transcripts (data not shown). We further examined their presence in the genome of *P. dumerilii*; *id3252* is present in full, whereas only 140 bp of *id13944* (834 bp in length) aligns with the genome (data not shown). This partial alignment suggests that *id13944* may undergo post-transcriptional processing or the genome assembly has not captured its entire sequence. Given that both transcripts exceed >200 bp in length and potentially lack coding ORFs, we suggest that *id3252* and *id13944* might be long non-coding RNAs (lncRNA). Hypothesizing that they could interact with other non-coding elements such as miRNAs, we used lncTAR (Li et al., 2015) to predict RNA-RNA interactions. For this analysis, we used miRNAs known from *P. dumerilii* (data obtained from the Arendt Lab). The results revealed that both *id13944* and *id3252* might interact with numerous microRNAs (miRNAs) identified in *P. dumerilii* (Supplementary Materials 9 and 10). These putative lncRNAs could be involved in regulating some of the tissue and sex-specific expression patterns we observed in this study and will be an interesting line of inquiry in future studies.

## Discussion

### Divergent transcriptional programs in spermatogenesis and oogenesis

This study unraveled the sex-biased gene expression patterns in the germline of the marine annelid *P. dumerilii*. The analysis of gene ontology (GO) enrichment for differentially expressed (DE) genes indicated distinct molecular programs underlying female and male gametogenesis. Male-biased genes were predominantly associated with spermatogenesis and related pathways, including “sperm principal piece”, “positive regulation of histone H3K36 methylation”, ”“sperm capacitation” and “CatSper complex” (Fig. 4A). This enrichment of sperm-related pathways in males corroborates previous findings on the transcriptional profiles of males other annelids (Álvarez-Campos et al., 2019; Ponz-Segrelles et al., 2020). Notably, components such as calcium ion binding and CatSper, a sperm-specific Ca2+ channel, are consistently related to spermatogenesis across metazoans (Castro-Arnau et al., 2022; Jiang et al., 2024; Ren et al., 2001; Speer et al., 2021; Wu et al., 2020). Calcium signaling and CatSper play crucial roles regulating sperm behavior, such as flagellar beating and sperm motility mechanisms in ascidians (Chung et al., 2017; Kijima et al., 2023; Ren et al., 2001). The enrichment of the gene ontology term "positive regulation of histone H3K36 methylation" suggests that the conserved regulatory mechanisms governing H3K36 trimethylation, which play a crucial role in the histone-to-protamine transition, are also active during spermatogenesis in *P. dumerilii* (Gardiner-Garden et al., 1998; Powers et al., 2016; Shirane et al., 2020). This suggests that our RNA-seq analysis captured transcriptional signatures associated with both early and late stages of spermatogenesis in *P. dumerilii* juveniles (Fig. 6D). The presence of these conserved spermatogenesis-related pathways, including CatSper function and histone H3K36 methylation, in the transcriptional profile of *P. dumerilii* gonial clusters underscores the shared molecular mechanisms of spermatogenesis across metazoans.

Conversely, the female-biased genes were primarily linked to metabolic processes and cellular components, enriched in categories such as “macromolecule metabolic process”, “organic substance metabolic process”, “organelle”, and “membrane-bounded organelle” (Fig. 4B). Enrichment in cellular components and metabolic processes have been observed in female gonads of vertebrates (Cao et al., 2023; Dong et al., 2021; Wang et al., 2017) and invertebrates (Hart and Foster, 2013; Peng et al., 2015), pointing to the oocyte maturation process. Female transcription profiles of other annelids have also indicated enrichment for cellular components (Ponz-Segrelles et al., 2020) and metabolic pathways (Álvarez-Campos et al., 2019). In *P. dumerilii*, oocytes at late stages undergo vitellogenesis characterized by oocyte enlargement and yolk production (Fischer, 1974; Fischer, 1975). Such cell growth requires higher metabolic activity and increased number of organelles in the ooplasm such as mitochondria, endoplasmic reticulum, and Golgi complexes (Gu et al., 2015; Van Blerkom, 2011). These findings highlight that the mechanisms thought to be conserved for gametogenesis are also present in the marine annelid *P. dumerilii*.

### Developmental stages and sex-biased gene expression

The analysis of differential expression and validation of DE genes showed significant sex-biased transcriptional profiles in 50S and 60S worms, with less differences in 40S worms. These results reflect the increased expansion of gonial clusters to the trunk and progression of gametogenesis stages in 50S and 60S (Kuehn et al., 2022; Meisel, 1990). The expansion of gonial clusters to the trunk typically occurs when worms are around 35-40S (Kuehn et al., 2022). At this initial phase of expansion, gonial clusters may be in mitotic stages (spermatogonia or oogonia), which morphologically resemble each other closely (Fischer, 1974; Fischer, 1975; Meisel, 1990). Our study showed less sex-biased expression of DE genes in the gonial clusters of 40S worms (Fig. 2B), indicating that at mitotic phase male and female gonial clusters exhibit similar transcriptional profiles. Alternatively, 40S worms, being smaller and typically possessing fewer gonial clusters during early expansion to the trunk (Kuehn et al., 2022), may yield samples more enriched for coelomocytes (somatic cell types found in the coelom along with gonial clusters). Even though some coelomocytes such as eleocytes are known to start showing sex–biased differences at later stages of juveniles such as production of vitellogenin in females (Fischer and Hoeger, 1993; Schenk and Hoeger, 2020), they may not have significantly different transcriptional profiles at this earlier stage.

On the other hand, 50S and 60S worms showed pronounced sex-biased expression of DE genes in the gonial clusters (Fig. 2B). Prior research indicated that 50S worms predominantly feature gonial clusters in mitosis, while 60S worms present advanced stages, including gonial clusters in early meiosis (Fischer, 1974; Meisel, 1990). Therefore, we expected to capture the mitotic and early meiotic stages of spermatogenesis with our RNAseq on 40S, 50S and 60S worms. However, we observed 60S worms with maturing sperm, which is a post-meiotic stage of spermatogenesis (Fig. 4A’-D’, Fig. 6A). Additionally, 14 out of 24 mature males had 55- 69 segments (Supplementary Material 1), indicating that 50S and 60S males may exhibit maturing gametes at post-meiotic stages at much earlier segment numbers than we originally expected. This indeed aligns with our results of GO enrichment for sperm-related genes in males, such as “sperm principal piece”, and “CatSper complex” (Fig. 3A).

In females, gonial clusters are mitotic at the onset of expansion to the trunk and later enter meiosis, which is arrested in Prophase I of meiosis until fertilization (Fischer, 1974; Nakama et al., 2017). Based on this, our RNA-seq data covered mitotic (oogonia) to early meiotic oocytes in 40-60S worms (Fig. 6E). Oogenesis occurs with no cell division after meiosis arrests, but oocytes undergo characteristic cell growth and vitellogenesis. Oogenesis in *P. dumerilii* has been characterized in four stages: 1. polygonal oogonia, 2. synaptic oocyte, 3. late prophase I oocyte and 4. vitellogenic oocyte, which might be dissociated from the cluster (Fischer, 1974). Interestingly, juveniles display asynchronous stages during early oogenesis, as a single individual may simultaneously present oocytes at different stages (Fischer, 1974). In agreement with the former studies, we observed the simultaneous presence of oogonia and oocytes at different stages in 50S and 60S worms (Fig. 4E-H, Fig. 5A), meaning early oogenesis is asynchronous. Oocytes become uniform in size as they approach a diameter ∼40µm (Fischer, 1974), after which they grow gradually and synchronously to reach a final size of ∼160µm (Fischer et al., 2010). This growth phase is usually characterized by an elevated metabolic turnover (Fischer and Hoeger, 1993), aligning with our findings on GO enrichment analysis, which showed enrichment for metabolic pathways in female-biased genes (Fig. 3B). Therefore, female-biased transcription patterns might be enriched as oogenesis advances into synaptic and vitellogenic phases.

Altogether, these findings indicate that the transcriptional transitions are detectable via bulk RNAseq, RT-PCRs and *in situ* HCRs from the 50S stage onwards, aligning with the increased expansion of gonial clusters to the trunk and their transition from mitotic to meiotic stages of gametogenesis.

### Comparison of the anterior cluster and gonial clusters

The anterior cluster, a group of germ cells located near the head, emerges from 4 primordial germ cells located adjacent to the PGZ during larval stages (Özpolat et al., 2017; Rebscher et al., 2007; Rebscher et al., 2012). These 4 primordial germ cells migrate towards the pharyngeal region, where they proliferate and persist throughout most juvenile stages (Kuehn et al., 2022; Özpolat et al., 2017; Rebscher et al., 2007; Rebscher et al., 2012). Visible in the pharyngeal region before and after the expansion of the gonial clusters to the trunk, the anterior cluster becomes indistinct and dispersed in later stages of gametogenesis, particularly when spermatocytes and vitellogenic oocytes become abundant in 60S and 70S worms (Fig. 5A, Supplementary Material 8) (Metzger and Özpolat, 2024). The morphology of the anterior cluster nuclei resembles that of spermatogonia and oogonia, featuring large and elongated nuclei (Figs. S5 and S6) (Rebscher et al., 2007). Experiments using Dil-labeling to track cells suggested that the anterior cluster as the source of gonial clusters (Rebscher et al., 2007).

Our results and analyses revealed an absence of sex-biased expression of DE genes in the anterior cluster despite anterior cluster being a component of the reproductive system (Fig. 6A-B, Fig. S7). Ultimately, there are no consistent sex-specific markers for the anterior cluster, irrespective of developmental stage (e.g. 40S, 50S, or 60S). However, our analysis is limited to the expression patterns of the selected genes. Further investigation, such as conducting RNAseq of the anterior cluster and comparing it with GCs at different developmental stages of juvenile males and females could reveal potential sex-biased transcriptional profiles in this tissue. As isolating the anterior cluster poses challenges, we propose that single-cell RNAseq holds promise for profiling the different cell types (anterior cluster, gonial clusters, and the growth zone -see below) and finding specific markers that can be used as cell-, sex- or stage-specific markers.

### Expression of DE genes in the posterior growth zone

Genes such as *vasa*, *piwi*, *nanos* are germline markers in many traditional research organisms such as the fruit fly, *C. elegans*, and zebrafish, however these same genes can display broader expression patterns, including in somatic tissues, multi- and pluripotent stem cells, and regeneration blastemas in other organisms such as annelids, echinoderms, and salamanders (Juliano et al., 2010; Özpolat and Bely, 2016; Poon et al., 2016; Zhu et al., 2012). This broader molecular program that is not confined to germ cells has been termed as the germline/multipotency program (Juliano et al., 2010). In *P. dumerilii* the germline/multipotency marker expression has been widely documented in the posterior growth zone (PGZ) and the regeneration blastema (Gazave et al., 2013; Metzger and Özpolat, 2024; Planques et al., 2019). To date, there has been no germ cell specific marker identified in *P. dumerilii*, although the PGZ stem cells specifically express *hox3* (Gazave et al., 2013). As our RNAseq data was collected from germ cell enriched tissues (and no PGZ), we asked whether the DE markers we identified for males and females were specific to gonial clusters and the anterior cluster (i.e germline-specific), or whether they were also expressed in the posterior growth zone (i.e. broader expression) similar to other germline/multipotency markers. We found that most of the male/female markers were present in the PGZ therefore generally there is no significant difference between PGZ, GCs, and the anterior cluster for the markers analyzed. We also did not find sex-biased expression differences of the selected markers in the PGZ (Fig. S7). However, two genes were more often absent or at very low expression levels in the PGZ: *lrrct* (3 out of 18 worms had expression), *kctd21* (3 out of 14 worms had expression). While these two genes are not completely absent from the PGZ in all worms analyzed, when present, they have very low expression in PGZ compared to gonial clusters, therefore they could be potentially good markers for the germ cells in the future.

### Sten exhibits a tissue-dependent sex-biased expression

In previous research, *sten* was identified as a male-biased gene in *P. dumerilii* in the brain samples of fully mature worms and worms that were at the edge of maturation (called pre-mature worms) (Schenk et al., 2019). Interestingly, in this study we found the opposite in the younger juvenile worms: *sten* is up-regulated in female gonial clusters, which demonstrates that a single gene can exhibit distinct expression patterns within the same individual, depending on the tissue, developmental stage, and sex. While *sten* is up-regulated in the male brain in pre-mature and mature individuals, it is also evidently required during oogenesis rather than spermatogenesis in younger juveniles of *P. dumerilii,* based on our RNAseq data and RT-PCRs. In addition, the expression of *sten* in the brain of the juvenile stages we focused on is similar between males and females, showing equally reduced expression compared to pre-mature and mature worms, as observed in our in situ HCRs, indicating that the expression differences in the brain possibly arise closer to maturation.

### dmrt1 is a male-biased gene in P. dumerilii gonial clusters

We identified four genes with DM domain in *P. dumerilii,* including *dmrt1, dmrt1/2*, *dmrt2*, and *dmrt3* (Fig. S8). Notably, *dmrt5* is not present in *P. dumerilii* and appears to have been lost in many annelids, except for the marine meiofaunal annelid *Dimorphilus gyrociliatus* (Ji et al., 2023) (Fig. S8). Among these groups of genes, only *dmrt1* was identified as a DE gene, being up-regulated in male gonial clusters.

*dmrt1* has been proposed as a sex determination gene in various animals (Picard et al., 2015). In fish, *dmrt1* has been found to be up-regulated in testes (Bar et al., 2016). In fruit flies, splice isoforms of DSX (another gene with DM domain) underlie sex differences, with male-specific isoforms activating genes and a female-specific isoform repressing them (Hildreth, 1965). In *P. dumerilii*, *dmrt1* is upregulated in both male brains (Schenk et al., 2019), and in male gonial clusters (this study), although also exhibiting lower expression levels in female gonial clusters (Fig. 6A). To investigate the potential involvement of spliced isoforms, we analyzed the expression of *dmrt1* isoforms. Our analysis revealed that females and males express all isoforms of *dmrt1*, but these are upregulated in males (Fig. S11). Hence, our data did not conclusively demonstrate differential isoform expression between males and females. Rather, it indicates a predominant bias towards males in expression levels.

However, our RT-PCR results indicate that *dmrt1* exhibits more pronounced sex-biased expression in 60S worms (advanced gametogenesis) compared to 40S worms (early gametogenesis). In line with our findings, *dmrt1* expression has been detected in undifferentiated gonads of male and female pufferfish, with a bias towards males (Yan et al., 2018). In pufferfish, expression levels of *dmrt1* are important in sex determination (Hu et al., 2019; Yan et al., 2018; Yan et al., 2021). Treatment with 17-beta estradiol reduces expression of *dmrt1*, leading to male-to-female sex change in pufferfish gonads (Hu et al., 2019). This pattern of increased expression of *dmrt1* in differentiated male gonads, while undifferentiated (bipotential) gonads show expression in both males and females is largely reported in the literature (Zarkower and Murphy, 2022). These transitions in *dmrt1* expression levels may be critical for the regulation of sexual determination and differentiation in *P. dumerilii* gametogenesis.

### A novel protostome-specific methyltransferase is associated with oogenesis

We identified a methyltransferase that is up-regulated in female gonial clusters, and potentially functions as a transmembrane methyltransferase in *P. dumerilii*. Our phylogenetic analysis revealed that this methyltransferase is closely associated with an “uncharacterized protein”’ found only in protostomes. This group of methyltransferases falls into a separate group from those characterized in recent phylogenies of methyltransferases (e.g. METTL1-27) (Wong and Eirin-Lopez, 2021). We termed it *psmt*, which stands for “protostome specific methyltransferase”, although our analysis has a predominantly spiralian coverage, as we selected the top hit sequences from BLAST annotation results (Fig. S14), nevertheless there were no similar genes found outside of protostomes.

Our analysis indicates that *P. dumerilii*’s *psmt* features the domain of S-adenosylmethionine (SAM) dependent methyltransferases. SAM-dependent methyltransferases belong to the Class I or Rossmann-like type of methyltransferases (Sun et al., 2021). This type of methyltransferase uses SAM as the methyl donor to methylate substrates (Martin and McMillan, 2002). Domain predictions from TMHMM and others, identified a transmembrane domain within *psmt’s* N-terminus (Figs. S16 and S17). Transmembrane domains usually show a function in facilitating the entry and exit of molecules, which is crucial for regulating several metabolic processes. This is consistent with our data which revealed an enrichment for metabolic processes in female gonial clusters in *P. dumerilii* (Fig. 3B). Therefore, *psmt* could be involved in metabolic processes in *P. dumerilii* female gonial clusters. Its specific presence in protostomes, specially in aquatic species (Fig. S14), potentially indicate that these mechanisms of methylation are conserved in protostome oogenesis.

### Sex-biased expression of long non-coding RNAs

We also detected and annotated putative long noncoding RNAs showing sex-biased expression in gonial clusters, hinting at their potential role in the genetic machinery of gametogenesis and sex-differentiation. *id13944* was identified as a putative lncRNA highly expressed in female gonial clusters, while *id3252* was highly expressed in male gonial clusters.

LncRNAs are known to play diverse regulatory roles in chromatin remodeling, epigenetic, transcriptional and posttranscriptional regulation (Mattick et al., 2023). Some lncRNAs are associated with sex-biased transcriptional differences mechanisms such as the X-inactive specific transcript (Xist) (Brown et al., 1991; Penny et al., 1996). Xist mediates X-chromosome inactivation in mammals via recruitment of the protein complex PRC2 (Silva et al., 2003; Zhao et al., 2008). This chromosomal dosage compensation is crucial for normal female development, which ensures the same dosage compensation as males with a single X chromosome (Engreitz et al., 2013; Gayen et al., 2016). However, the lncRNAs found in this study are shorter and not analogous to Xist (data not shown). This suggests that alternative regulatory mechanisms may be at play. For instance, many lncRNAs seem to play a role in sex differentiation, interacting with miRNAs in many animals (Dong et al., 2021; Feng et al., 2021; Mysore et al., 2021; Yan et al., 2021; Zhang et al., 2018). lncRNAs can act as a decoy for miRNA that binds to specific mRNA targets, leading to increased expression of the mRNA targets (Ebert and Sharp, 2010). In the annelid *Eisenia fetida*, *neev*, a lncRNA has been related to the process of chaetogenesis acting as miRNA decoy (Patel et al., 2020). We identified numerous miRNAs that might interact with both *id13944* and *id3252* lncRNAs in *P. dumerilii* (Supplementary Materials 9 and 10), hinting at a role as a miRNA decoy. The precise functions of these lncRNAs remain to be tested in future studies.

### Insights into sex determination mechanisms in P. dumerilii

Sex determination systems are categorized into two types: genetic determination, which can be chromosomal or polygenetic, and environmental determination, which is influenced by environmental cues, such as temperature (Chadwick and Goode, 2002; Crews, 2003; Piferrer, 2021). However, in many cases, a combination of both genetic and environmental factors may contribute to sex determination (Piferrer, 2021). During evolutionary transitions from gonochorism (separate sexes) to simultaneous hermaphroditism (individuals producing both male and female gametes simultaneously), presence of intermediate stages consisting of species exhibiting environmental sex determination or sequential hermaphroditism has been proposed (Leonard, 2018; Weeks et al., 2006). This phenotypic plasticity in intermediate evolutionary stages tends to be governed by epigenetic mechanisms (Piferrer, 2021).

*P. dumerilii* is a gonochoric species, meaning each individual will produce either female or male gametes (Fischer et al., 2010). However, *P. dumerilii* also lacks heterochromosomes (Jha et al., 1995) and the existence of sex *loci* involved in genetic sex determination remains uncertain. On rare occasions, we have observed oocytes in *P. dumerilii* males (data not shown), which could point towards a more plastic nature of sex determination in *P. dumerilii,* such as epigenetic regulation. Moreover, given that its sister species *P. massiliensis* is a sequential hermaphroditic species (Lücht and Pfannenstiel, 1989), we suggest that the gonochorism in *P. dumerilii* may be a recent/ intermediate evolutionary trait in the group.

This study does not resolve the mechanisms governing sex determination, but offers some evidence of an epigenetic program governing sex differentiation in *P. dumerilii*, such as: i) the presence of different methylation programs in males (e.g. enriched histone H3K36 methylation pathways) and the up-regulation of a specific methyltransferase in females (*psmt*); ii) The relatively late appearance of sex-biased transcription profiles in 50S worms may suggest that epigenetic changes take place during or before this developmental stage, determining the fate of the germ cells; iii) the similarity of transcription profiles in 40S worms, and change in *dmrt1* expression biased to males in later stages (50S, 60S) may evidence bipotential germ cells in early stages. The precise genetic and molecular mechanisms underlying sex determination in *P. dumerilii* remain to be uncovered through further research.

## Conclusion

Our study generated a valuable resource for germ cell research in *P. dumerilii*. Using RNAseq, we characterized the transcription profiles of mitotic and meiotic germ cells in gonial clusters in *P. dumerilii*, capturing their transition to differentiated gametes.

Additionally, we provide data on the expression of sex-biased genes not only in the targeted gonial clusters but also in other tissues of the germ cell lineage and growth zone. Notably, sex-biased expression is detected in gonial clusters, while other germline tissues do not exhibit such differences, suggesting that these transcriptional biases appear during the expansion of gonial clusters to the trunk, which is potentially the developmental stage where sex determination occurs. Our study uncovered *dmrt1*, a conserved gene found in the double-sex mab-3 domain, as a male-biased gene associated with gametogenesis. Importantly, we characterized novel genes such as a protostome-specific methyltransferase, *psmt*, and long non-coding RNAs with sex-biased expression which is expressed in gonial clusters tending to meiotic stages.

These findings highlight that both conserved molecular programs might govern gametogenesis in *P. dumerilii*. Additional comparative studies are required to enhance our comprehension of the evolutionary foundations and mechanisms governing this biological phenomenon.

## Methods

### Animal cultures

Juvenile worms were maintained following our lab’s standard protocol, ensuring consistent conditions of feeding, photoperiod and temperature (Kuehn et al., 2019). However, the source of water varied based on the culture site. At the Marine Biological Laboratory, we used 1 µm-filtered natural seawater (salinity ranged from 32–34 parts per thousand (ppT), at the Washington University in St. Louis, cultures were maintained in artificial seawater (ASW, salinity of 35 ppT), prepared with distilled water and red sea salt (Red Sea Fish).

The sex ratio was monitored for the entire culture of worms, with collections conducted at least twice a week from August 2021 to February 2024. On each collection day, males and females are separated from cultures. Maturing worms that have not completed maturation are set aside as “unknown”.

### Sample Collection for Bulk RNAseq

We divided the worms into three groups based on their age or size, measured by the number of segments: **40S**, 40-49 segment-long; **50S**, 50-59 segment-long; and **60S**, 60- 69 segment-long. These groups are representative of the course of gametogenesis, reflecting both early mitotic (40-49 segment-long) and meiotic stages (50 segment-long or longer) (Fig. 1B-D).

In *P. dumerilii*, germ cells form gonial clusters, which are clusters of germ cells surrounded by sheath cells. These gonial clusters are located within the coelomic cavity of *P. dumerilii*. In addition to gonial clusters, the coelom also contains eleocytes. Due to the challenge of separating these cell types and considering the potential for eleocytes to also exhibit sex-biased transcriptomes, we opted to collect the coelomic contents (eleocytes and gonial clusters) for RNAseq. The coelomic contents were collected from amputated posterior bodies individually, eliminating the need to sacrifice the worms.

This allowed us to keep their anterior bodies alive to regenerate and mature, so the sex could be assigned to each sample.

For collection of coelomic contents, the worms were relaxed in 1:1 mixture of 7.5% MgCl2 diluted in filtered artificial sea water (ASW) for 15-30 min prior to dissection. Subsequently, the worms were bisected with scalpels at the midbody. The bisection site was determined based on the worm’s body length, after the chaetigerous segment number ∼15-20 (Fig. 1A, Supplementary Material 1). Next, the posterior halves were specifically dissected in filtered sea water (FSW) using sterilized surgical material (scalpel and needles) designed to exclusively isolate the coelomic contents and avoid contamination from the gut (ÖzpolatLab-GitHub-Ribeiro, 2024). The resulting sample volume consisted of approximately 50-100 μL of coelomic contents mixed with ASW. These samples were used for RNA isolation.

The anterior body (head fragments) of each worm was cultured individually in 50 mL plastic cups until maturation. Worms were fed 0.335 mL of 2.0 g/L spirulina (Micro Ingredients) and 0.3 g/L Sera micron flakes (Sera, 0072041678), twice a week. When found alive, mature worms were fixed for HCRs. After maturation, we were able to sex 43 samples, because two worms died during the experiment.

### Experiment for testing sex reversal upon regeneration

We selected 37 juveniles bearing 40-59 segments for this experiment, which typically have many gonial clusters across the trunk. The worms were relaxed for 15-30 min in a 1:1 solution of 7.5% MgCl2 and 35 ppT ASW. After relaxation, the worms were bisected either after the chaetigerous segment 10, or at the midbody, removing about 20-30 posterior segments (Supplementary Material 2). The amputated posterior bodies were fixed for HCR *in situ* hybridization, while the anterior bodies were cultured to regenerate and grow at least 50 segments. Once worms reach approximately 50 segments, their gonial clusters undergo morphological changes following gametogenesis stages.

Therefore, we raised the worms to about 50 segments (2-3 months after amputation) and fixed them. Whole-mount HCR was performed with all worms using a combination of gene probes targeting *vasa* (positive control for gonial cluster), *dmrt1* (male marker) and *psmt* (female marker). To determine sex reversal, we compared gonial clusters before and after regeneration for each individual, assessing the morphology of the gonial clusters and expression of the selected markers.

### RNA isolation

mRNA was isolated from samples of coelomic contents collected as explained above. Samples were then centrifuged at 3000 g for 2 minutes. Supernatant was removed and cell pellets were immediately processed for RNA extraction using the PicoPure kit following the provided instructions for cell pellets (Life Technologies, KIT0204), including a step of DNase treatment with Qiagen 79254. Total RNA was measured on a Thermo Scientific Nanopore 2000. Samples with RNA concentrations measuring less than 30 ng/μL were excluded from sequencing or RT-PCRs.

### RNA sequencing

Samples of RNAseq were kept in -80L no longer than two months and sent for sequencing at the Genome Technology Access Center, McDonnell Genome Institute, Washington University in St. Louis. The facility prepared the libraries using mRNA-Seq cDNA Amplification - Clontech SMARTer. Libraries underwent quality check on a 2100 Bioanalyzer (Agilent Technologies) and samples were paired-end sequenced on one lane of an Illumina NovaSeq S4 2x150 instrument. The obtained reads were uploaded to NCBI Sequence Read Archive (SRA) (Supplementary Material 1).

### Transcriptome assembly

After sequencing, raw reads of the 45 samples sequenced were quality checked using FastQC v.0.11.5 (http://bioinformatics.babraham.ac.uk/projects/fastqc/), and Trimmomatic v0.38 (Bolger et al., 2014) was used to trim adapters and filter low quality reads with the options ILLUMINACLIP HEADCROP:10 LEADING:34 TRAILING:30 MINLEN:36. Next, reads were submitted to quality check again using FASTQC (https://www.bioinformatics.babraham.ac.uk/projects/fastqc/) and the results were analyzed with MultiQC (Ewels et al., 2016). Quality control results are available in a repository associated with this study (ÖzpolatLab-GitHub-Ribeiro, 2024).

For the generation of a reference transcriptome assembly to use in our analyses, reads of 18 samples were selected with three replicates for each condition of sex and age (Supplementary Material 1). These reads were then concatenated into two files as they were paired-end sequences. These files were used as input -left and -right for de novo assembly using Trinity v2.14.0 (Grabherr et al., 2011; Grabherr et al., 2013; Haas et al., 2013). To obtain transcriptome statistics of the assembled transcriptome, we ran the TrinityStats.pl script of Trinity v2.14.0. Assembly and transcriptomic statistics were conducted at the RIS Scientific Compute Platform of the Washington University in St.

Louis (https://docs.ris.wustl.edu). In addition, we ran a completeness analysis against the metazoa_odb10 database using a dockerized BUSCO v5.3.2 (ezlabgva/busco:v5.3.2_cv1) (Simão et al., 2015). Both software were operated within a custom Docker image (https://hub.docker.com/r/rannyele/transcriptomics).

### Functional annotation and gene ontology

We annotated the transcripts using the Trinotate pipeline (https://github.com/Trinotate/Trinotate.github.io/wiki) (Bryant et al., 2017). First, we identified coding regions and predicted coding sequences with TransDecoder v5.5.0 (Haas, BJ. https://transdecoder.github.io/) containerized in Docker (quay.io/biocontainers/transdecoder:5.5.0--pl5321hdfd78af_5). Subsequently, both the assembled transcripts and the predicted coding sequences were annotated. For annotation, we employed BLAST+ (Altschul et al., 1990; Camacho et al., 2009), HMMER (http://hmmer.org/) (Eddy, 2011), signalP v6 (Teufel et al., 2022), DeepTMHMM (Hallgren et al., 2022), Infernal (Nawrocki and Eddy, 2013), gene ontology (GO) (Gaudet et al., 2017), EggnogMapper (Cantalapiedra et al., 2021), and DIAMOND (Buchfink et al., 2021), using Trinotate v.4.0.0 via Docker (trinityrnaseq/trinotate). The annotation report was generated with Trinotate.

### Differential expression

To find differentially expressed (DE) genes, we conducted both a preliminary and an in-depth analysis. In the pilot analysis, we compared only the 18 samples used for the transcriptome assembly, all from worms that had matured first in our experiment and passed quality control. Using bowtie2 (Langmead et al., 2009) for transcript quantification and DEseq (Love et al., 2014) in R, we obtained a preliminary list of differentially expressed transcripts. As more worms matured and samples could be labeled, we expanded the number of replicates for the differential expression analysis. Although we generated RNAseq data for 45 samples, we selected 30 samples for differential expression to ensure an equal representation of sexes and sizes, with 5 replicated per condition. Trinity (v.2.8) Differential Expression module was used to compare expression profiles at the isoform and gene (“Trinity gene”) levels. Transcript abundance for each sample was estimated using kallisto (Bray et al., 2016), while DESeq2 (Love et al., 2014) was employed for differential expression analyses.

Subsequently, those differentially expressed (DE) genes or isoforms that are at least 4- fold differentially expressed at a significance of p > 0.001 in any of the pairwise sample comparisons were extracted. This study focused on gene-level results, but isoform-level results were also performed. Trinotate annotation was integrated to provide the Gene Ontology categories (GO) terms enriched or depleted in these DE genes. GOseq (Young et al., 2010) was employed to visualize the results of gene ontology enrichment analyses. The results and outputs on these analyses are available in our repository (ÖzpolatLab-GitHub-Ribeiro, 2024).

### Identification of candidate genes

We used our preliminary differential expression analysis to identify sex-related candidate genes, in addition to BLASTn searches of sex-related genes documented in the literature (Picard et al., 2021; Schenk et al., 2019) within our transcriptome assembly. The identified genes were cross-referenced with the list of DE transcripts from the preliminary DE analysis (Supplementary Materials 3-6). The BLAST searches led to the selection of two DE genes candidates for sex differentiation: *dmrt1* and *sten* (Supplementary Material 4). Due to a large report of spliced isoforms of dmrt-like genes as important for sex differences in animals (Picard et al., 2015), we retrieved the transcript sequences for all its six predicted isoforms of *dmrt1* to analyze expression patterns across samples, which were visualized in heatmaps.

Subsequently, we filtered our list based on the lowest adjusted p-value (FDR, false discovery rate), ranging from 3.1E-45 to 3.21E-10 (Supplementary Materials 5 and 6). This refined list was used to select 12 additional genes in the top50 rank of more significant p-values. Notably, these selected genes largely corresponded to genes ranked within the top 200 DE genes revealed by the in-depth differential expression analysis, except two of them: a zfp-like gene (TRINITY_DN514_c3_g1) and the transcript TRINITY_DN36_c4_g3_i1 (Supplementary Material 3). These two genes were then removed from the final list of DE candidate genes for validation, which contained a total of 12 genes, being 10 selected from the top50 DE genes, and two genes from literature (Supplementary Material 3).

### RT-PCRs

The validation of the selected DE genes was performed using reverse transcription polymerase chain reactions (RT-PCRs) (Supplementary Material 7). Primers were designed based on the sequences of the transcripts obtained from our transcriptome assembly, using the NCBI primer blast tool (https://www.ncbi.nlm.nih.gov/tools/primer-blast/) (Supplementary Material 12). mRNA samples were obtained following the methods described above (RNA isolation). The total isolated mRNA was reverse transcribed using First Strand cDNA Synthesis Kit for RT-PCR (Sigma Aldrich 11483188001). RT-PCRs were conducted using Taq polymerase (Invitrogen EP0401). Amplicons were analyzed using 1% agarose gels in Tris-(TBE).

### Hybridization chain reaction (HCR) probe design

We designed HCR split-probes for selected sex-related genes revealed by our RNAseq using our custom software (Kuehn et al., 2022). The software generates pairs of 25 bp sequences that are complementary to the target DNA sequence, compatible with selected HCR™ amplifiers from Molecular Instruments, Inc (https://www.molecularinstruments.com/). We selected up to 32 pairs of oligo pools per gene or combined genes. The custom probes were synthesized to an amount of 5 nmol per DNA oligo pool byIntegrated DNA Technologies (https://www.idtdna.com) and re- suspended to 50 µM with Nuclease Free water (Invitrogen UltraPureTM Distilled Water, 10977-015). Targeting 14 genes of interest, we tested 14 probe sets and 6 of them were successful for the following genes: *dmrt1*, *psmt*, *kctd21*, *id625*, *lrrct*, *sten*, *id3252* and *id13944* (Suppl. Mat. 12).

### Tissue collection and whole-mount HCR

Whole worms were collected for HCRs, anesthetized in 1:1 0.22um FNSW: 7.5% MgCl2 for 15-30 minutes and fixed overnight in 4% paraformaldehyde (PFA) diluted in 2X phosphate saline solution (PBS) and filtered artificial sea water (ASW). Fixed material was dehydrated in a gradient of 25%, 50%, 75% and 100% methanol series, up to 10 min each incubation step. Methanol solutions were diluted in 1X PBS with 0.1% Tween (1X PBSt). PBS solutions were prepared with diethyl pyrocarbonate treated ultrafiltered water. After dehydration, samples were stored in 100% Methanol at -20L until use.

HCRs were performed following previous protocols (Kuehn et al., 2022), with a modification in the form of substituting the proteinase K treatment with a Tween-20- based permeabilizing solution (Bruce et al., 2021). The probe hybridization buffers, wash buffers, amplification buffers and HCR™ amplifiers were obtained from Molecular Instruments, Inc (https://www.molecularinstruments.com/). Samples were mounted on cover slides with SlowFade™ Glass Soft-set Antifade Mountant, with DAPI (Invitrogen™ S36920) or SouthernBiotech™ Fluoromount-G™ Slide Mounting Medium (SouthernBiotech™ 010001).

### Statistics

To quantify RT-PCR products, agarose gels were photo documented with a LI-COR D-digit gel scanner. Gel bands were measured using Gel analyzer in Image J2 (version 2.14.0/1.54f). The values were normalized to arbitrary ratios, using a gene-to- housekeeping gene ratio.

We assessed the expression of selected DE genes within the gonial clusters, anterior cluster, and posterior growth zone. Expression presence was evaluated using HCR images, recording both the presence and absence of gene expression. We adopted a very conservative approach, considering expression positive if the gene was detected in at least two sites within the analyzed structure (data summarized in Suppl.Supplementary Fig. S7 and Supplementary Material 8). Oocyte singlets were treated as part of the gonial cluster when present. Gonial clusters were imaged and evaluated based on the expression of vasa as a positive control, with the number of gonial clusters determined by the presence of vasa expression. The data of presence and absence was used to generate contingency tables, of presence and absence of expression for males and females. The contingency tables served as input to analyze male to female odds ratio. Contingency tables containing values equal to zero were normalized by addition of 1 unit to each entry, following the concept of the Haldane-Anscombe correction. A chi-square test was implemented to identify significant differences. Statistical differences were analyzed using a chi-square test for independence. We calculated the percentage of expression of the selected genes based on the total number of imaged gonial clusters of females and males, including standard deviation.

We calculated the sex ratio for the cultures in our lab using the cumulative sum of the total number of males and females collected from August 2021 until February 2024. A cumulative ratio of the difference between females and males was also performed. A chi-square test was conducted to assess significant differences between the number of males and females in our cultures over time.

### Gonial cluster staining

Gonial clusters were stained with phalloidin and DAPI to visualize nuclei and cell morphology, facilitating the establishment of morphological parameters for male and female identification based on previously established literature (Fischer, 1974; Fischer, 1975; Meisel, 1990). Staining was performed in freshly collected gonial clusters using methods detailed above (see Sample collection for Bulk RNAseq). The gonial clusters were fixed 60 min on ice, in 4% PFA made with 16% paraformaldehyde, 2X PBS, and 0.22um NBSS in a ratio of 1:2:1. They were washed 3x with 1X PBSt, 5 min each wash; followed by staining with 1:100 DAPI (10ug/uL) (D9542, Sigma-Aldrich) and 1:40 phalloidin (A12379, Invitrogen) for 45 min in the dark at room temperature. Before imaging, samples were rinsed once with 1X PBSt.

### Imaging and visualization

Live images of mature worms were captured using a Zeiss scope Stemi 305 coupled with a built-in camera and an iPad 7th generation, software version 15.3.1, using the software Zeiss Lab Scope (version 4.2). Laser scanning confocal images were acquired with a Zeiss LSM 900. Images were analyzed and processed using FIJI ImageJ2 (version 2.14.0/1.54f) (Schindelin et al., 2012). All figures of this manuscript were edited using Adobe Illustrator, version 26.3.1.

### Phylogenetic analyses

The following approach was used for building all gene trees for *dmrt*, *psmt*, and *SoxB*. The *in-silico*-translated product of these genes were used to identify other similar *Platynereis dumerilii* transcripts on the Jekely Lab BLAST portal (https://jekelylab.ex.ac.uk/blast/). All libraries (*Platynereis dumerilii* MPI ESTs, *Platynereis dumerilii* transcriptome assembly version 1, *Platynereis dumerilii* transcriptome assembly version 2) were searched with BLAST and all hits were collected. These nucleotide sequences were *in silico* translated using Expasy’s Translate function (https://web.expasy.org/translate). The protein sequences used to produce each gene tree were also collected from previous literature (Heenan et al., 2016; Jurkowski and Jeltsch, 2011; Wang et al., 2023; Wexler et al., 2014; Wong and Eirin-Lopez, 2021) and combined with the *Platynereis* sequences. SeqKit v2.1.0 was used to remove duplicate sequences (Shen et al., 2016).

To improve the alignment of the proteins, the full length sequences were passed to InterProScan (https://prosite.expasy.org/scanprosite) to identify their respective family domains. The protein sequence of these domains were aligned using MAFFT (https://mafft.cbrc.jp/) with “allow gappy alignment” and “Auto” model choice (Katoh et al., 2019). Sequences were then pruned by first filtering with CD-HIT at 100% identity (Li and Godzik, 2006) , and then the MaxAlign function (Gouveia-Oliveira et al., 2007), ensuring that the *Platynereis* and Outgroup sequences were retained, then realigning these suggested sequences. The resulting MAFFT alignment was then passed to IQTree2 (Minh et al., 2020; Nguyen et al., 2015) for the tree search and ultrafast bootstrapping, SH-aLRT, and approximate Bayesian analyses. Trees were plotted with iTOL (https://itol.embl.de/) (Letunic and Bork) and then were edited in Adobe Illustrator CC 2024.

### Protein characterization and AlphaFold2 structure prediction

Modeling DMRT1 - The *Platynereis* DMRT1 protein was modeled as a monomer and as a dimer using ColabFold v1.5.5 (Mirdita et al., 2022), a web portal providing access to AlphaFold2 (Jumper et al., 2021). The program was run with settings of “num-relax: 1” and “template_mode: pdb100” (Jumper et al., 2021; Mirdita et al., 2022; Steinegger and Söding, 2017). ColabFold’s top ranked model was imported into ChimeraX v1.7.1 and, using the “Matchmaker” function, was aligned to the PDB crystal structure 4YJ0.

*psmt* AlphaFold2 models were created as different homo-oligmeric forms (monomer, dimer, trimer, tetramer, pentamer, hexamer, and octamer) using ColabFold v1.5.5 (Mirdita et al., 2022). With the exception of the octamer, the program was run with settings of “num-relax: 1” and “template_mode: pdb100”; the octamer was run with the same settings except “num-relax” was set to “0”. Irrespective of oligomeric state, the 50 amino acids of the models’ N-terminus had weak confidence (lDDT score < 60) (Jumper et al., 2021; Mirdita et al., 2022; Steinegger and Söding, 2017). Further *psmt* domain analysis was investigated using SMART (http://smart.embl-heidelberg.de), InterProScan (https://prosite.expasy.org/scanprosite), and TMHMM-2.0 (https://services.healthtech.dtu.dk/services/TMHMM-2.0). ColabFold’s top model was imported into ChimeraX v1.7.1 and superimposed upon crystal models 3BUS and 3E23 pulled from PDB via the “Matchmaker” function.

### Predicted protein structure visualization

*“*Molecular graphics and analyses performed with UCSF ChimeraX, developed by the Resource for Biocomputing, Visualization, and Informatics at the University of California, San Francisco, with support from National Institutes of Health R01-GM129325 and the Office of Cyber Infrastructure and Computational Biology, National Institute of Allergy and Infectious Diseases” (Goddard et al., 2018; Meng et al., 2023; Pettersen et al., 2021).

### AI tools

This study benefited from revisions of code and grammar using ChatGPT and Perplexity (OpenAI, 2023; Perplexity AI, 2023).

## ACKNOWLEDGEMENTS

We thank Özpolat Lab members, WashU Germ Cell Interest Group and Kerry Kornfeld Lab for their helpful feedback on this project. We thank Bria Metzger and Julianna Escudero for the gonial cluster images in Figure 1B-D, and our undergraduate students Kaitlyn Hong and Lisa Cheung for help with RT-PCRs.

## DATA AVAILABILITY

Coding resources, protocols, analyzed data, inducing data not shown, from this study can be accessed through the GitHub repository page https://github.com/BDuyguOzpolat/Ribeiro-et-al_Sex_Differentiation (ÖzpolatLab-GitHub-Ribeiro, 2024). Imaging data including light and laser scanning microscopy are available at Zenodo doi:10.5281/zenodo.11619572 (Ribeiro et al., 2024). RNAseq raw reads are available at SRA/NCBI (XXXXXX).

---

**Supplementary Figure S1.**
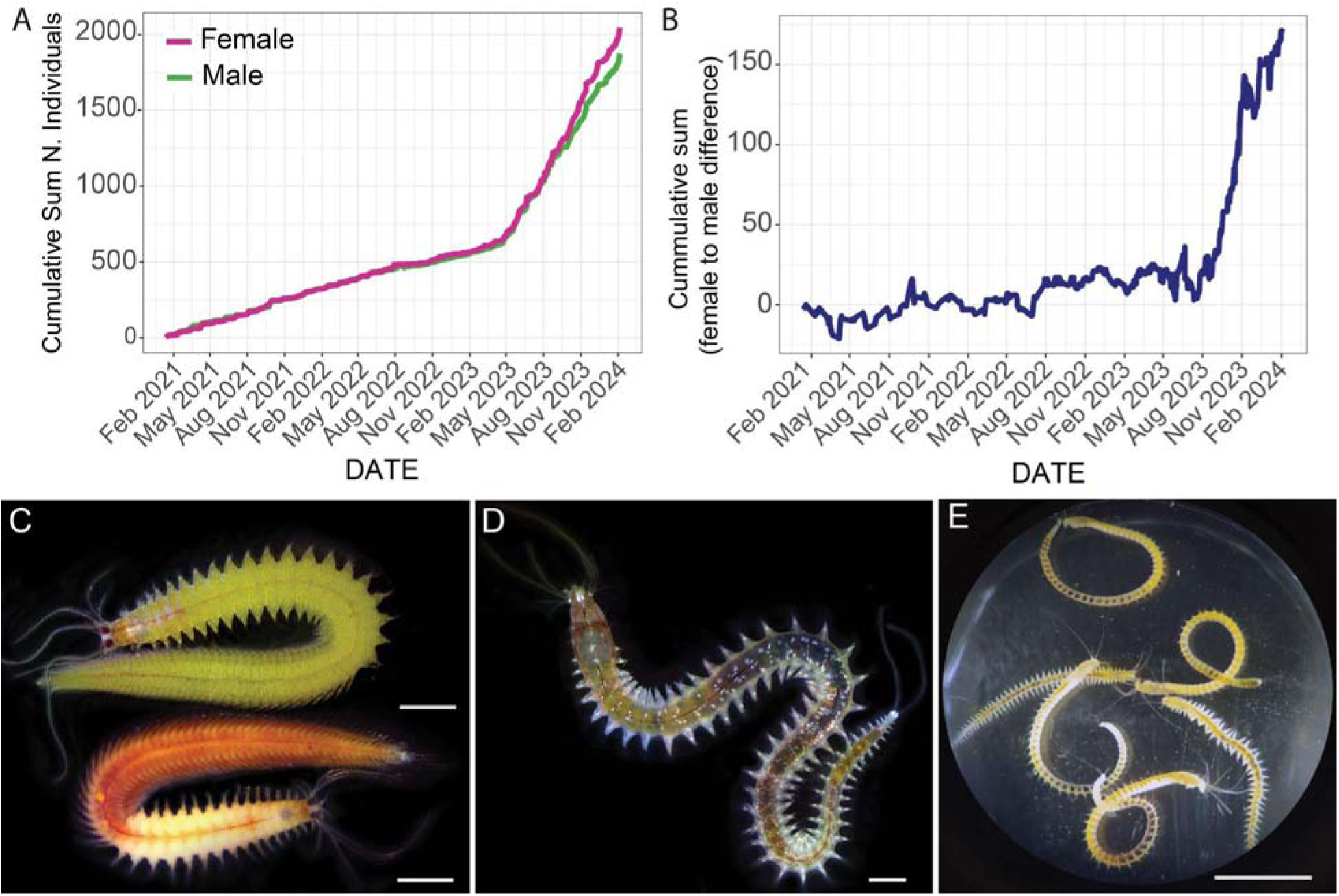
Male and female distribution in cultures and external morphology of mature and juvenile *P. dumerilii*. (A) Cumulative sum of male and females collected in the Özpolat lab cultures, considering only wild type worms. No differences are seen in the collection of males and females from February 2021 until August 2023, as the curves of males and females are overlapping. B) Female to male cumulative difference in lab cultures over three years (2021-2024). From February 2021 to August 2023, the female-to-male ratio in the lab cultures was approximately 1:1, with a near zero cumulative difference between the two genders. Starting in November 2023, there was an increase in the number of females compared to males. This change can be attributed to two factors: 1) The expansion of cultures in August 2023 increased the overall number of worms collected, leading to a steeper increase in the cumulative difference; 2) From August 2023 onwards, the involvement of trainees in identifying mature worms introduced a bias in counting females, while many males have been counted as premature (unknown sex). (C) Mature worms have sexual dimorphism, with males appearing red and white, while females are yellow. Other morphological traits differentiate males from females, such as the modified tail with papillae in males and a body constriction named as the epitokous border. While the epitokous border in males is typically after the segment number 15 (±1), the epitokous border in females can be after segment number 22 (±2) (Schulz et al 1989). (D) Juvenile image of a worm with 52 segments. Juveniles also present a different morphology when compared with mature worms, differing in the type of chaetae, size of eyes and swimming behavior but juveniles are similar to each other. (E) Examples of many juveniles with identical morphology, including males and females.

**Supplementary Figure S2.**
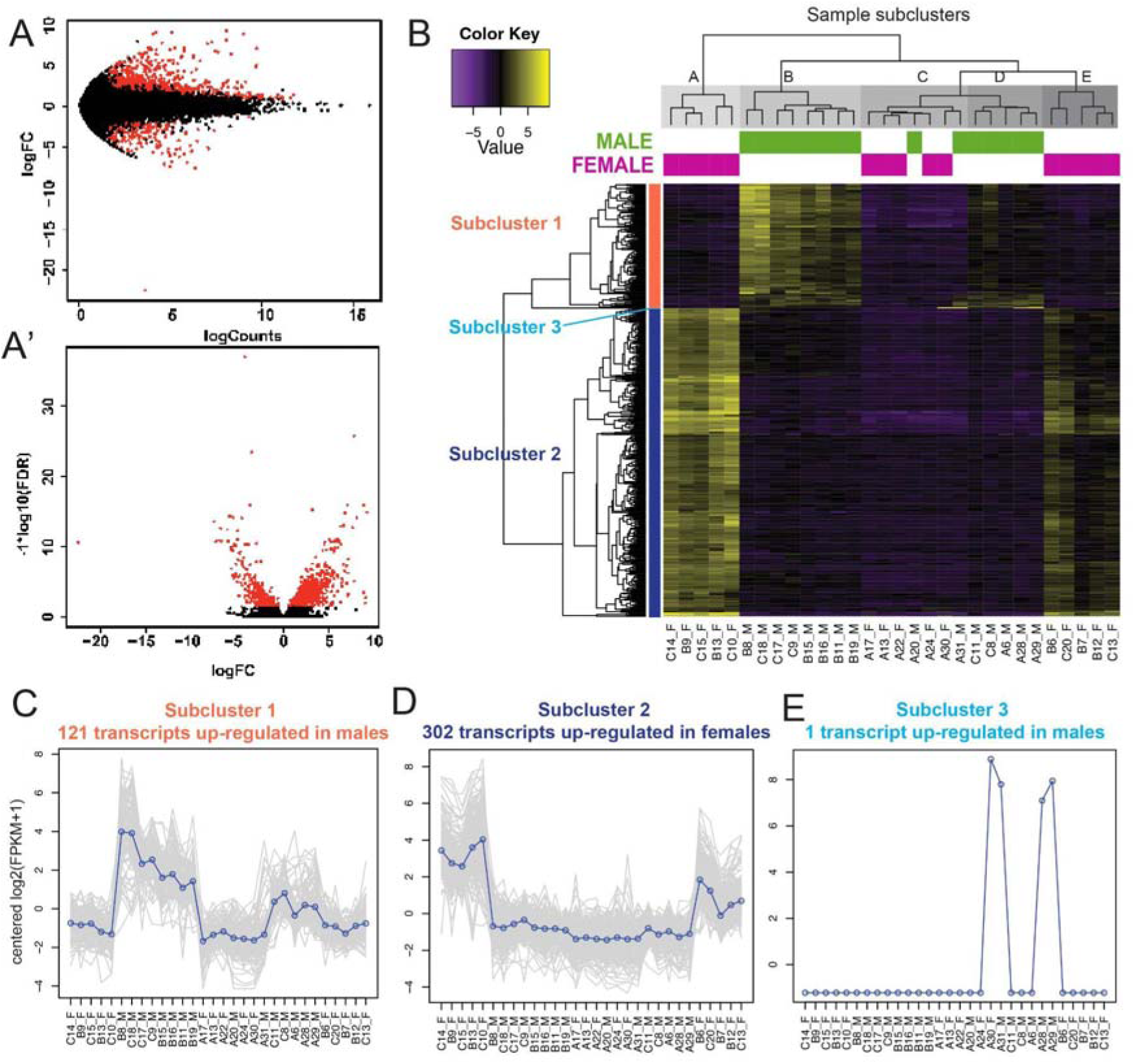
Sex-biased gene expression in *P. dumerilii* juveniles, additional details. (A-A’) Results of the differential gene expression analysis at the level of Trinity genes. (A) MA plot shows the distributions of Trinity ‘genes’ according to their log-to-fold change (logFC), adjusted p value <0.01. (A’) Volcano plot shows a false discovery rate (FDR) ranging from 2 to 35, adjusted p value <0.01. (B) Heatmap of differentially expressed genes over 30 samples, 5 replicates for each sex and group (40S, 50S and 60S). Samples were clustered by similarity in gene expression patterns. There are two groups of females (subclusters I and V) and two groups of males (subclusters II and IV), while males and females of 40S, the younger group of worms group, cluster together (subcluster III). (C-E) Subclusters 1 and 2 of differentially expressed (DE) genes. (C) Subcluster 1 shows genes up-regulated in males. (D) Subcluster 2 includes all DEgenes up-regulated in females. (E) A third subcluster containing one gene up-regulated in males was generated by the analysis.

**Supplementary Figure S3:**
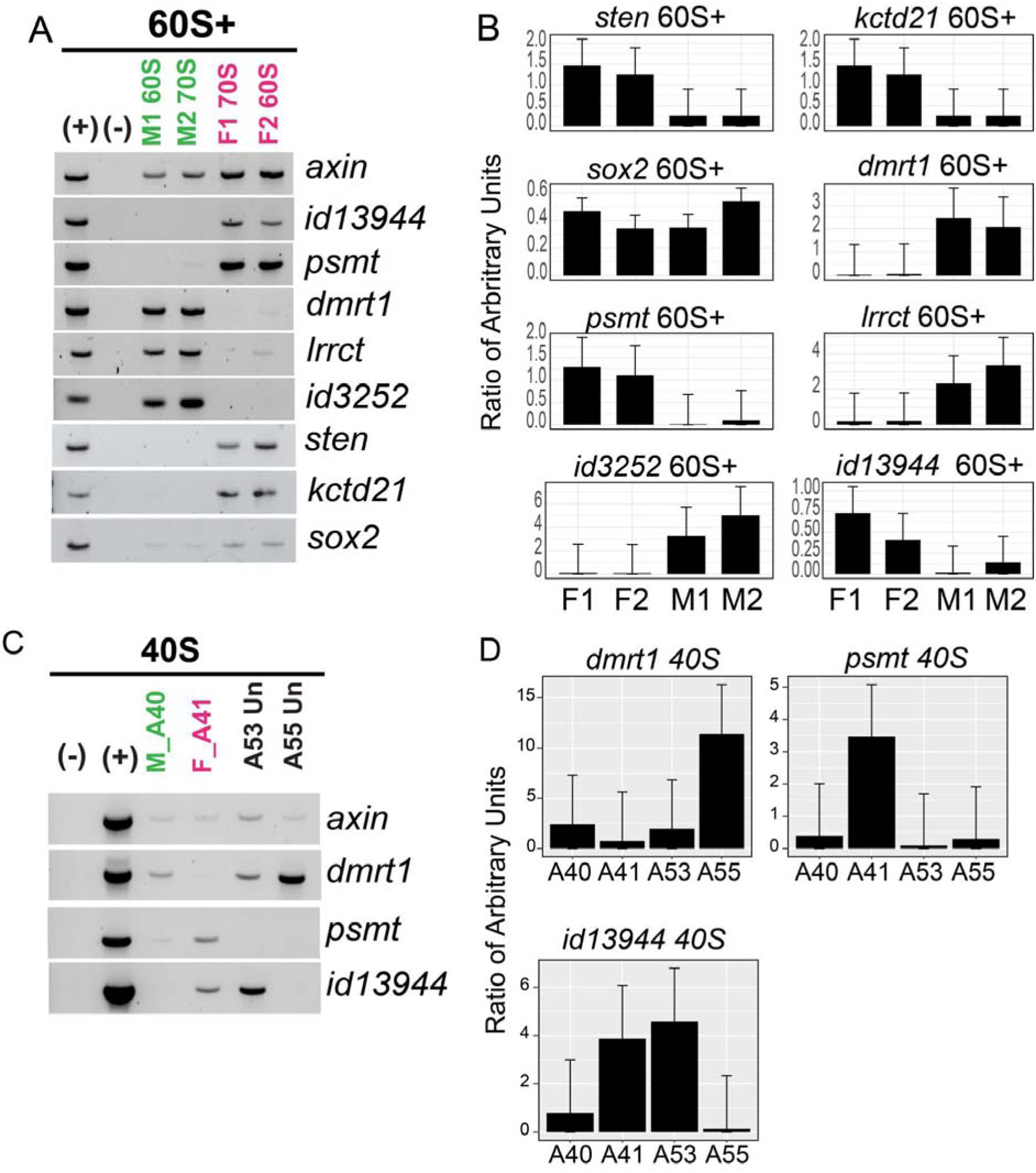
RT-PCR results and quantification of amplicons. (A) RT-PCR results for 8 genes in samples from males and females with 60-70S (B) Quantification of the results shown in A. (C) RT-PCRs on samples from 40S. A40 is male and A41 is female. The sex of A53 and A55 could not be determined based on RT-PCR results. (D) Quantification of gene expression from results shown in C. Quantification of *id10224* was not possible due to absence of positive samples (*axin*) for its agarose gel. Ratios of arbitrary units were calculated using the estimated expression of *axin* (housekeeping gene).

**Supplementary Fig. S4.**
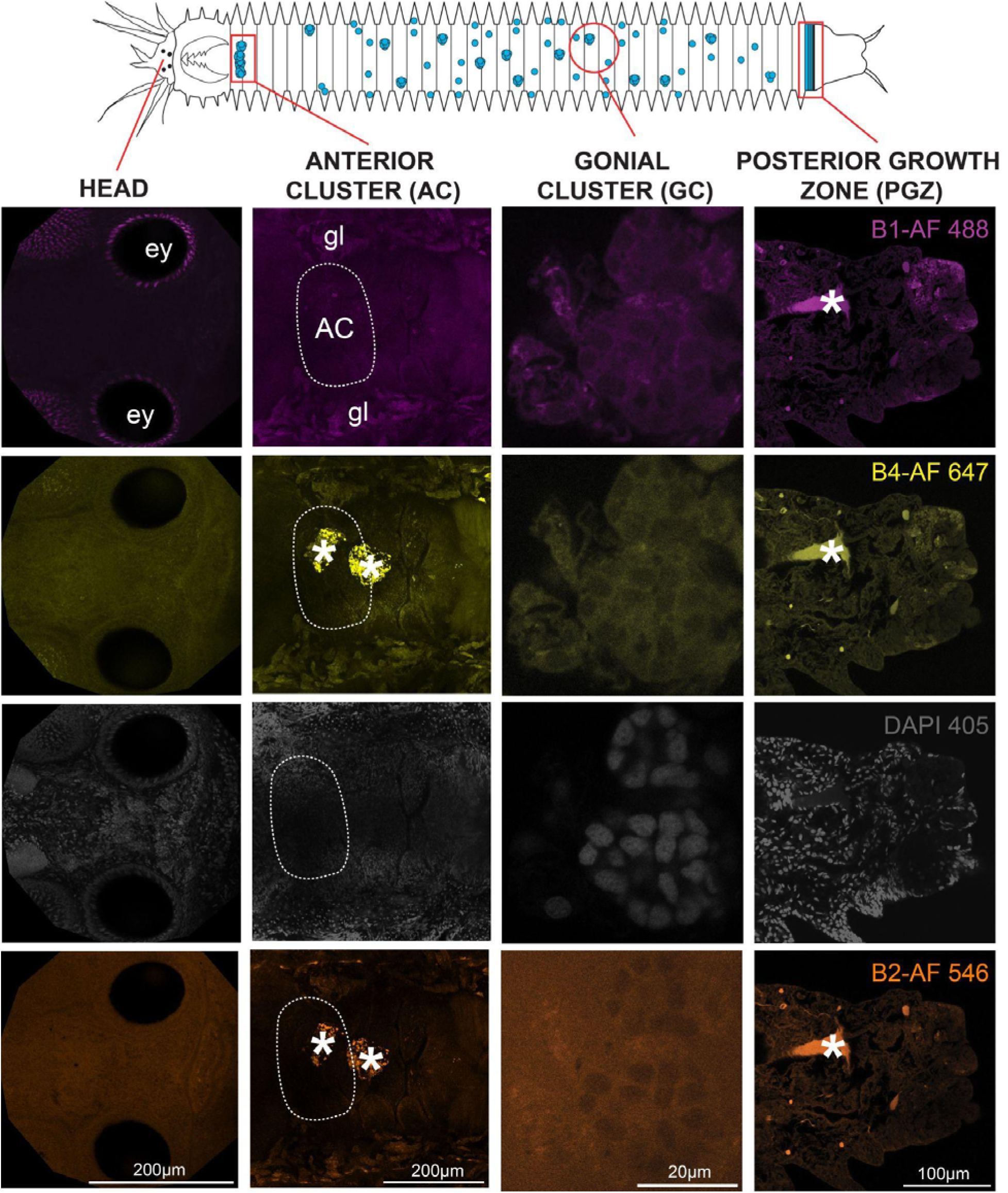
No-probe control whole-mount HCR images showing the head, the anterior cluster (AC), gonial clusters (GC), and posterior growth zone (PGZ). Images were taken with settings for Alexa fluorophores 488, 546, and 647, as well as DAPI (405). Dashed rectangles highlight the AC region, often showing background from the gut and accessory glands. To minimize background noise, AC images are zoomed in to focus on germ cells to analyze presence or absence of gene expression. Asterisks indicate gut content background. Orientation of all figures: anterior body to the left. Abbreviations: ey - eye, gl - glands, AC - anterior cluster.

**Supplementary Fig. S5.**
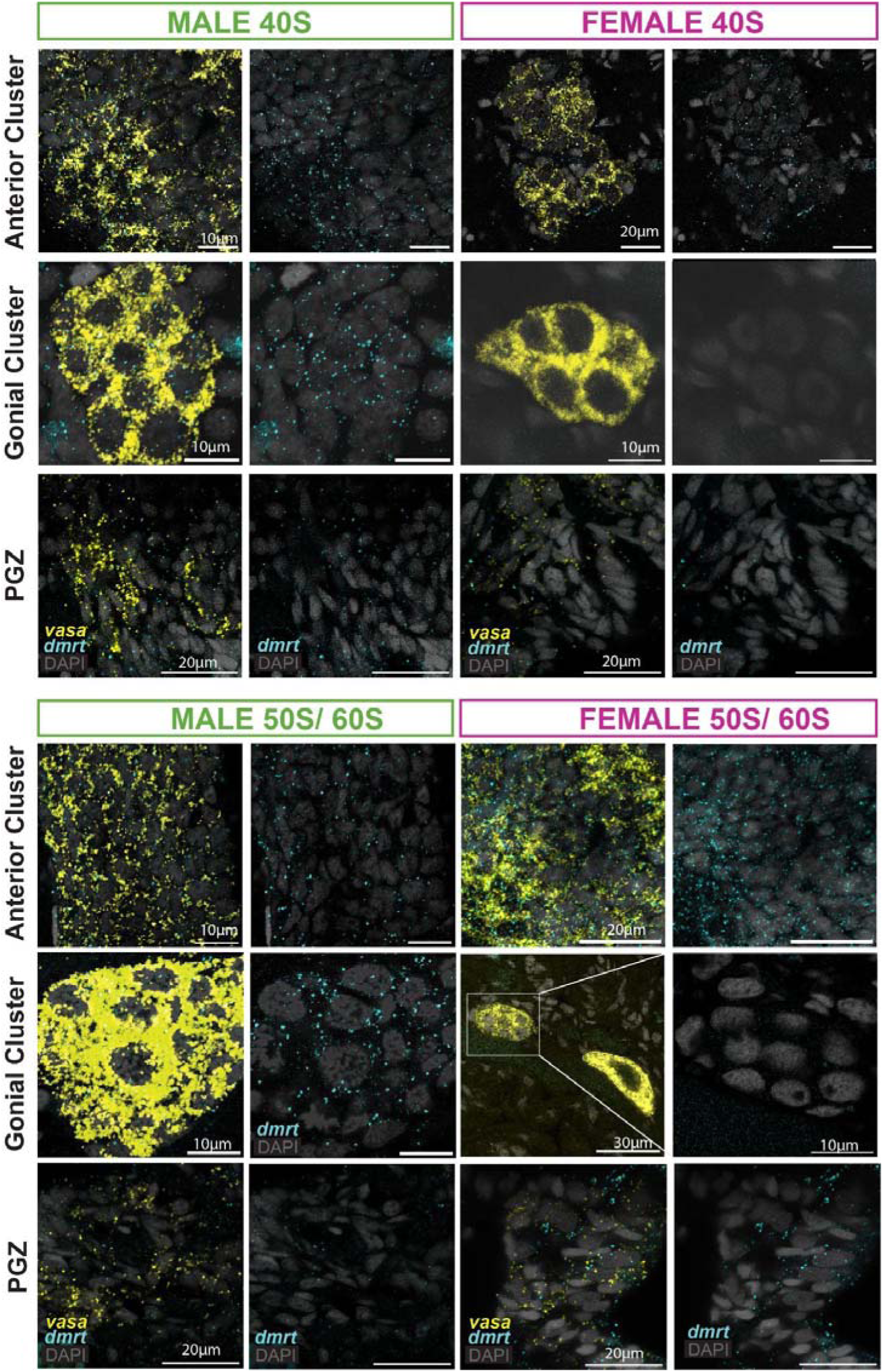
*dmrt1* sex-biased expression is limited to gonial clusters. Representative images of main results; both males and females express *dmrt1* in the anterior cluster and PGZ. In the gonial clusters, expression of *dmrt1* is biased towards males. Number of worms analyzed: 40S, males = 4, females = 1; 50S+ males n=5. 50S+ females = 8. Imaged 40S unknown worms, n= 19.

**Supplementary Figure S6.**
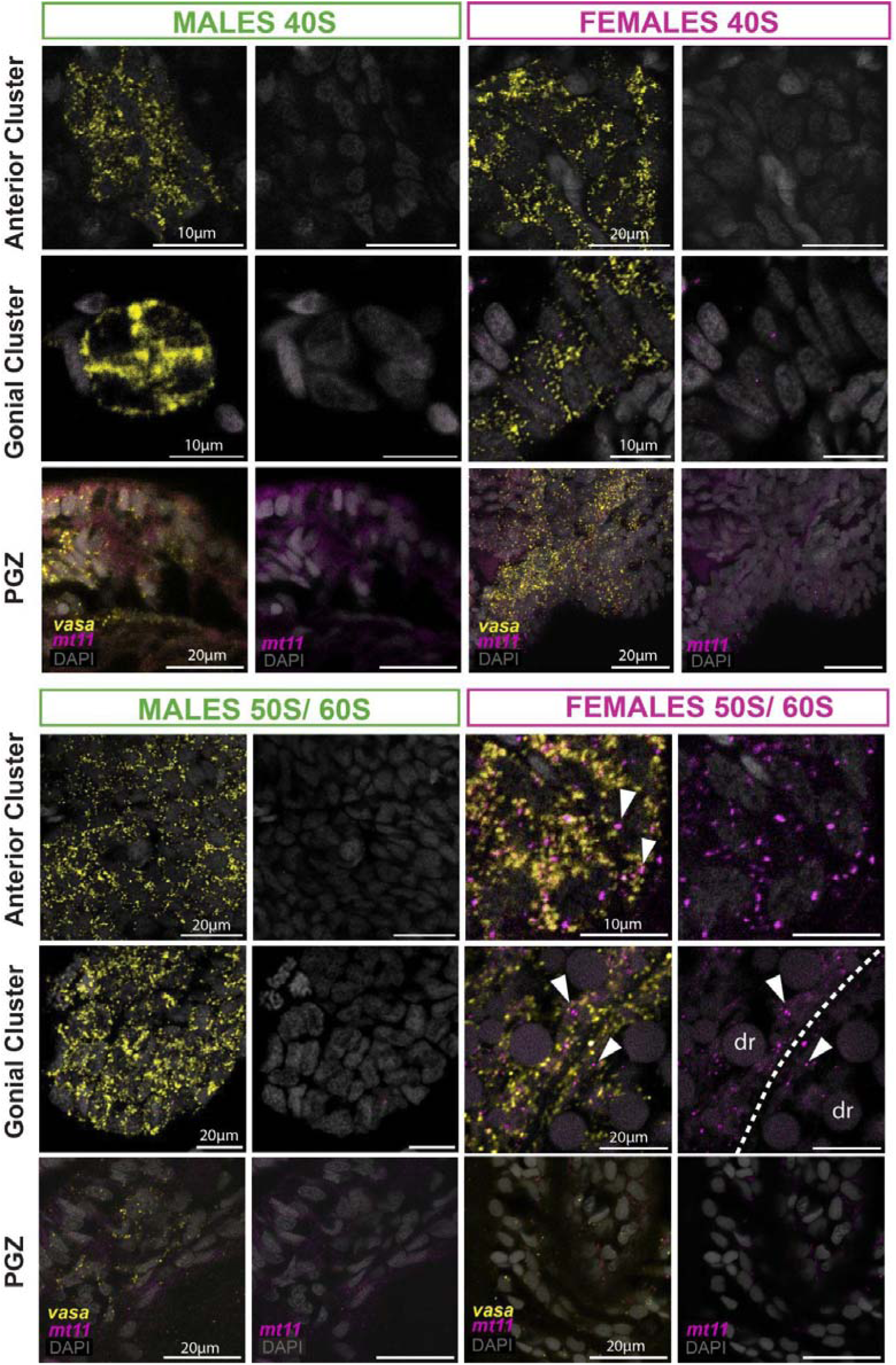
*psmt* expression patterns. *psmt* is highly expressed in females, particularly near the membranes of vitellogenic oocytes in 50S and 60S females (white arrowheads). Dashed lines mark the borders between oocytes, and arrowheads indicate sites of *psmt* expression near the cellular membrane. Number of analyzed worms: 40S, male = 2, female = 4; 50S+, male = 5, female = 8.

**Supplementary Fig. S7.**
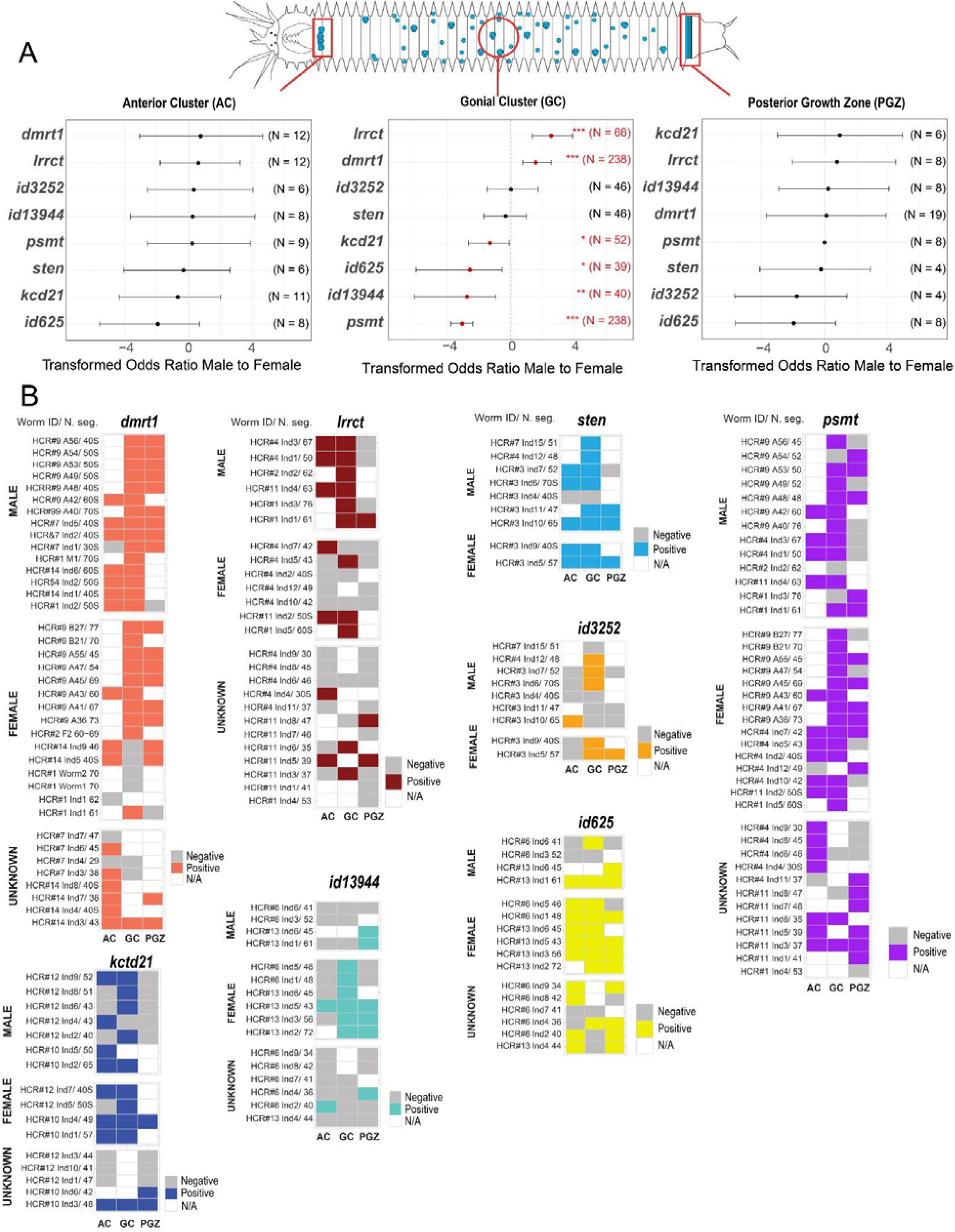
Summary of expression patterns of DE genes. (A) Odds ratios comparing male-to-female ratios of positive/negative gene expression in the anterior cluster (AC), gonial clusters (GC), and posterior growth zone (PGZ). (B) Graphical tables summarizing the expression status of each differentially expressed (DE) gene for each individual. "Positive" indicates the presence of gene expression, while "negative" indicates its absence. Expression was conservatively classified as positive if it was detected at very low levels in at least two sites. "N/A" denotes the absence of a given structure (AC, GC, PGZ). For example, young worms (20- 40S) that have not expanded the GC to the trunk, older worms (60S) where the AC is indistinguishable, and worms lacking a tail (absent PGZ).

**Supplementary Fig. S8.**
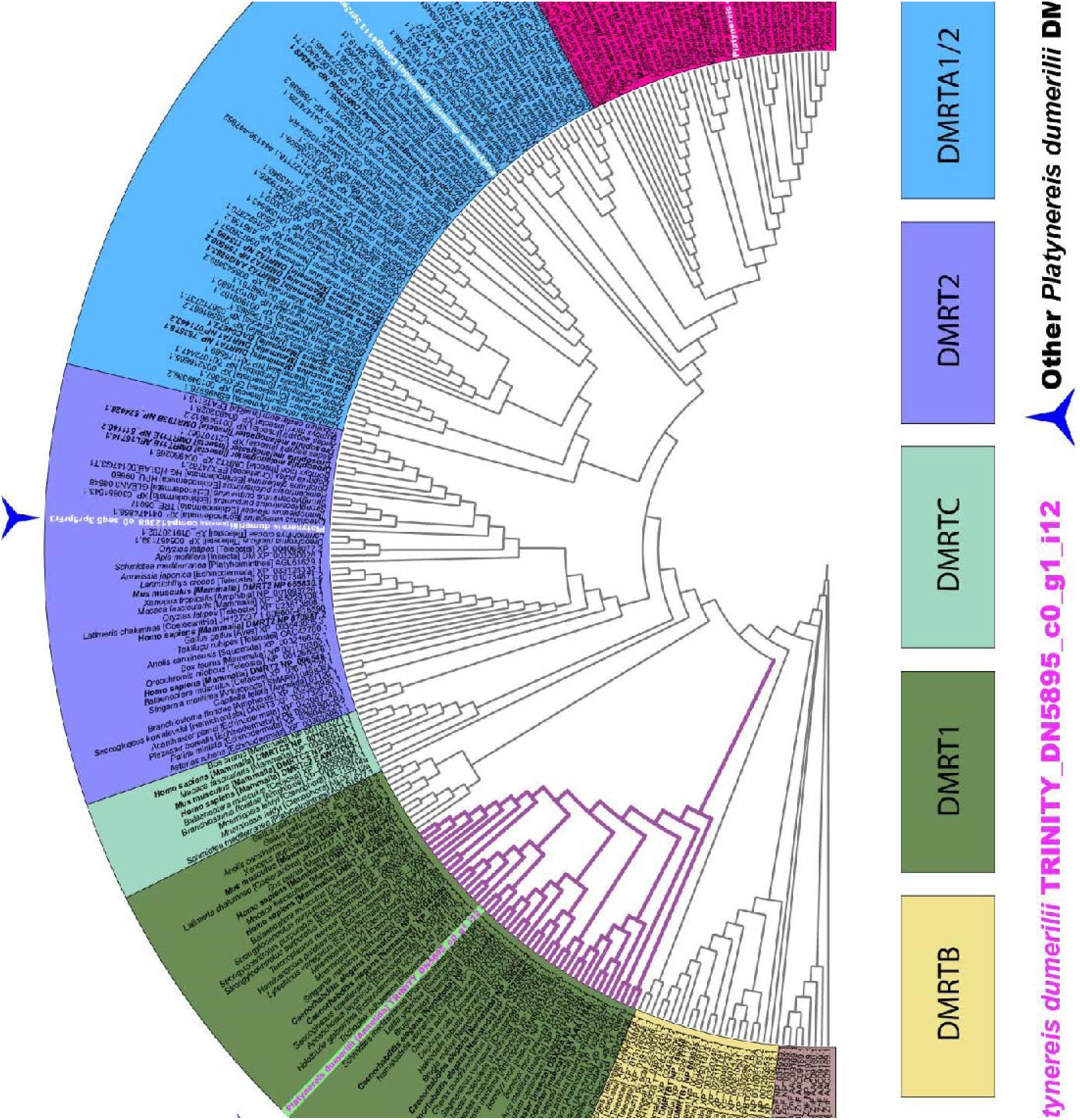
Analysis of *P. dumerilii* DMRT1 suggests functional DM domain. Approximate maximum likelihood tree of DM domains from DM-domain-containing proteins obtained from IQTree2. Protein assignment, indicated by background coloration, determined by co-clustering human, *Drosophila*, and mouse proteins (bold face IDs). The blue stars at the perimeter indicate *Platynereis dumerilii* sequences with the 5-pointed star showing the location of the DMRT1 sequence (from transcript TRINITY_DN5895_c0_g1_i12, a differentially expressed transcript) and the three-pointed stars indicating other transcripts falling within the DMRT gene family. The purple-stroked portion of the tree is further analyzed in Figs. S9 and S10. A high-resolution version of this tree can be found at the github repository (ÖzpolatLab-GitHub-Ribeiro, 2024).

**Supplementary Fig. S9.**
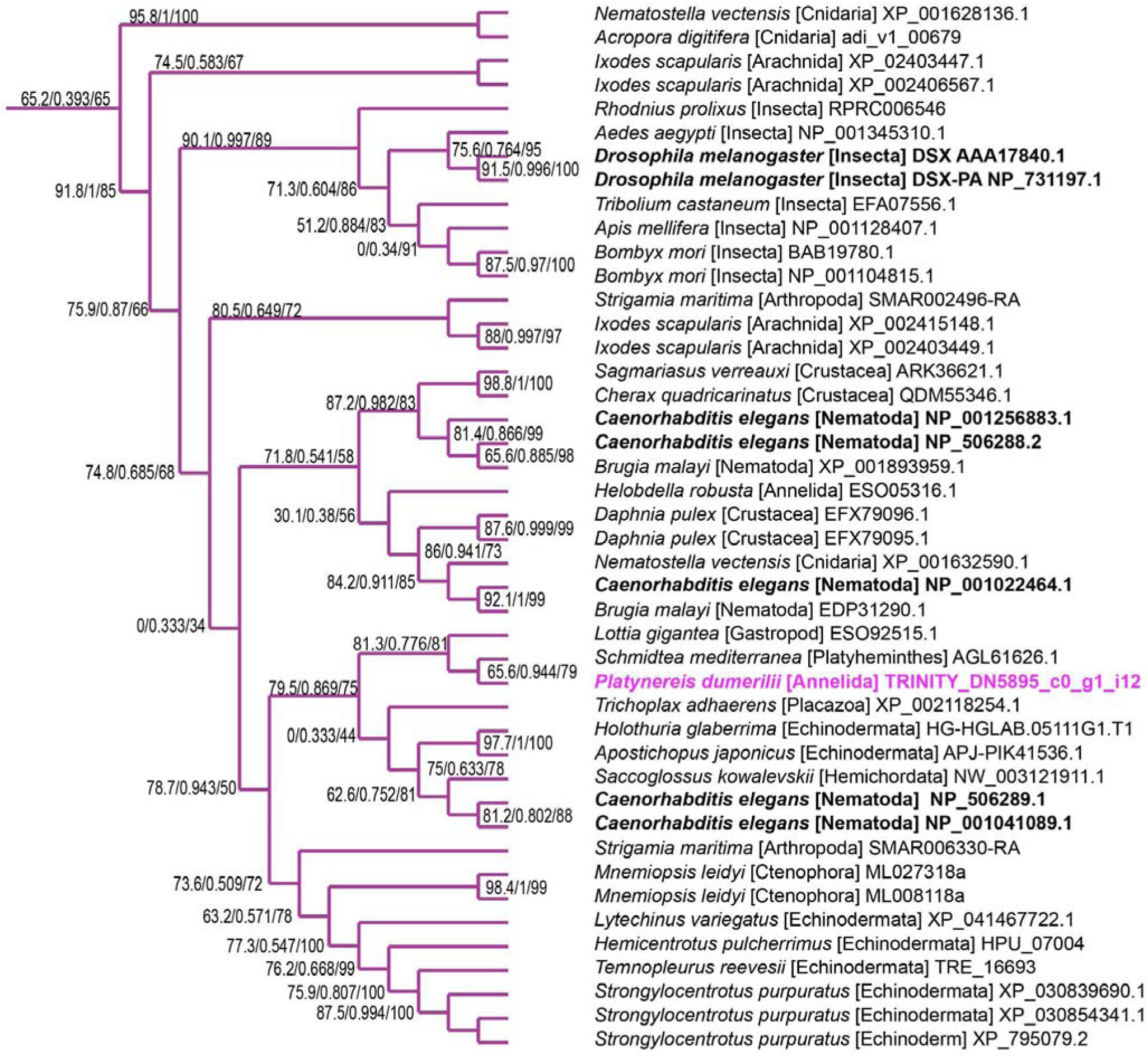
Subtree containing the *Platynereis dumerilii* DMRT1 domain. Support values are SH-alrt / Approximate Bayesian / UltraFastBootstrap as assigned by IQTree2.

**Supplementary Fig. S10.**
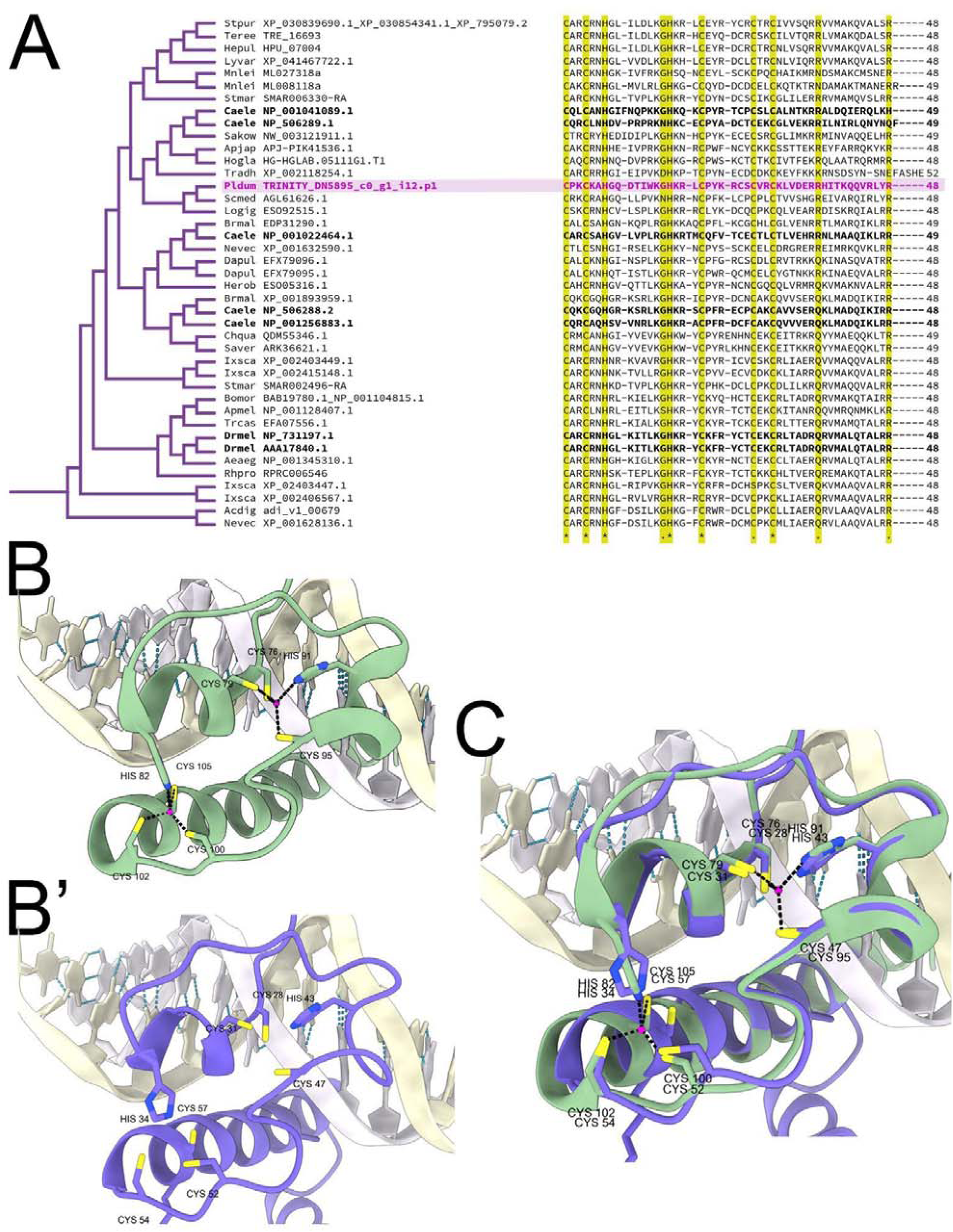
*P. dumerilii* DMRT1. (A) Sequence alignment of DM domains. *P. dumerilii* DMRT1’s DM domain shown with pink background. Conserved residues are indicated in yellow columns. (B, B’, C) ChimeraX renders a DM containing proteins on DNA. (B) Render of PDB 4YJ0’s crystal structure: Human DMRT1 DM domain (green) binding to DNA 25-mer (cream and white) showing the zinc (pink) and the zinc-binding Cysteine and Histidine residues (C76, C79, H82, H91, C95, C100, C102, and C105). (B’) Render of ColabFold’s top model of the entire *P. dumerilii* DMRT1 sequence spatially-aligned to 4YJ0 seen in B. Analogous Cysteine and Histidine residues (C28, C31, H34, H43, C47, C52, C54, and C57) are shown. (C) Overlap of Human and *P. dumerilii* DM domains structures bound to DNA.

**Supplementary Figure S11:**
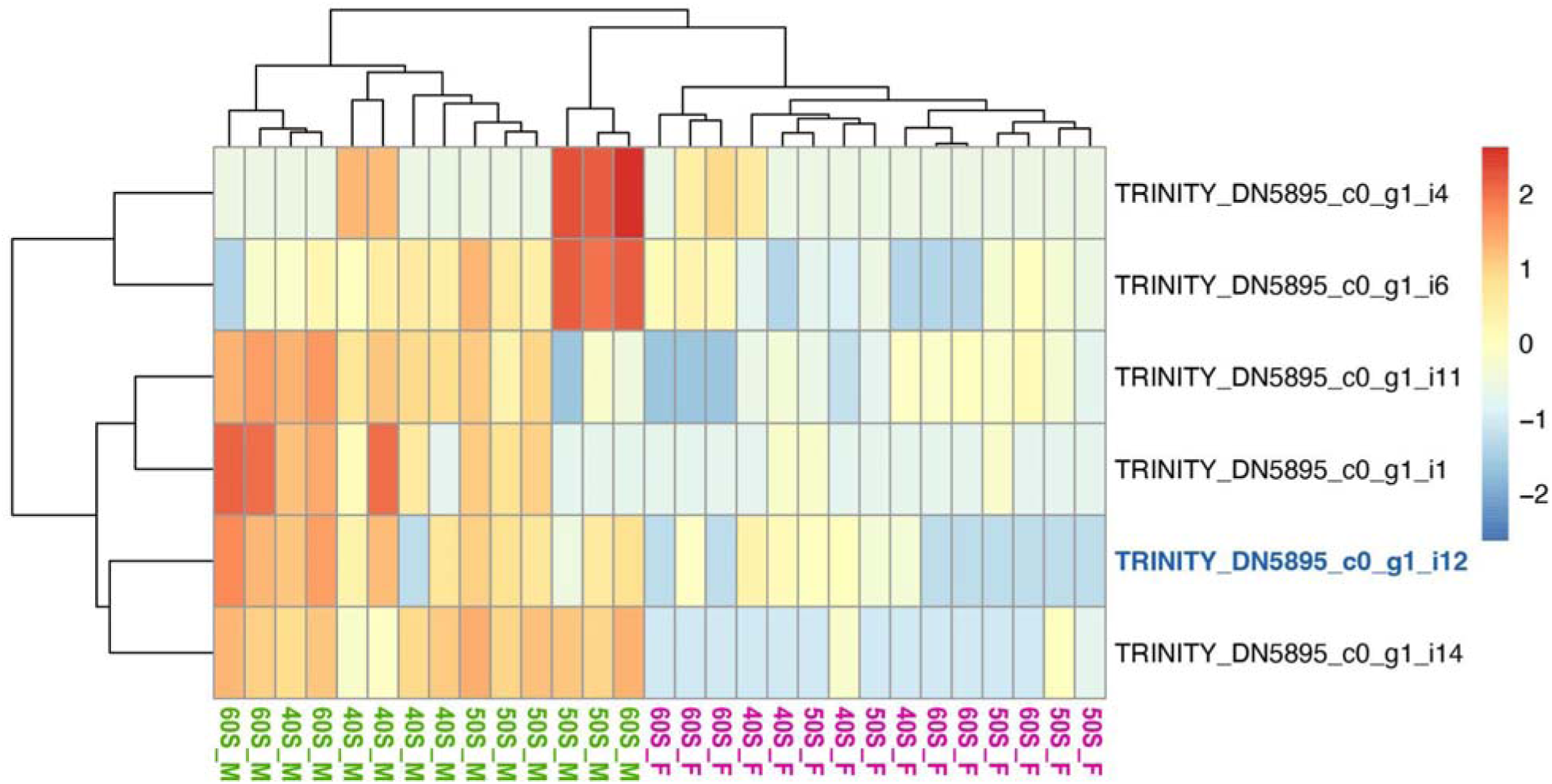
*dmrt1* isoform expression across males and females of *P*. *dumerilii*, from RNAseq analysis. Sample labels indicate developmental stage (40S, 50S and 60S) and sex (M, male; F, female). The transcript highlighted in blue was used to design oligo probes for *in situ* HCRs of *dmrt1*.

**Supplementary Figure S12:**
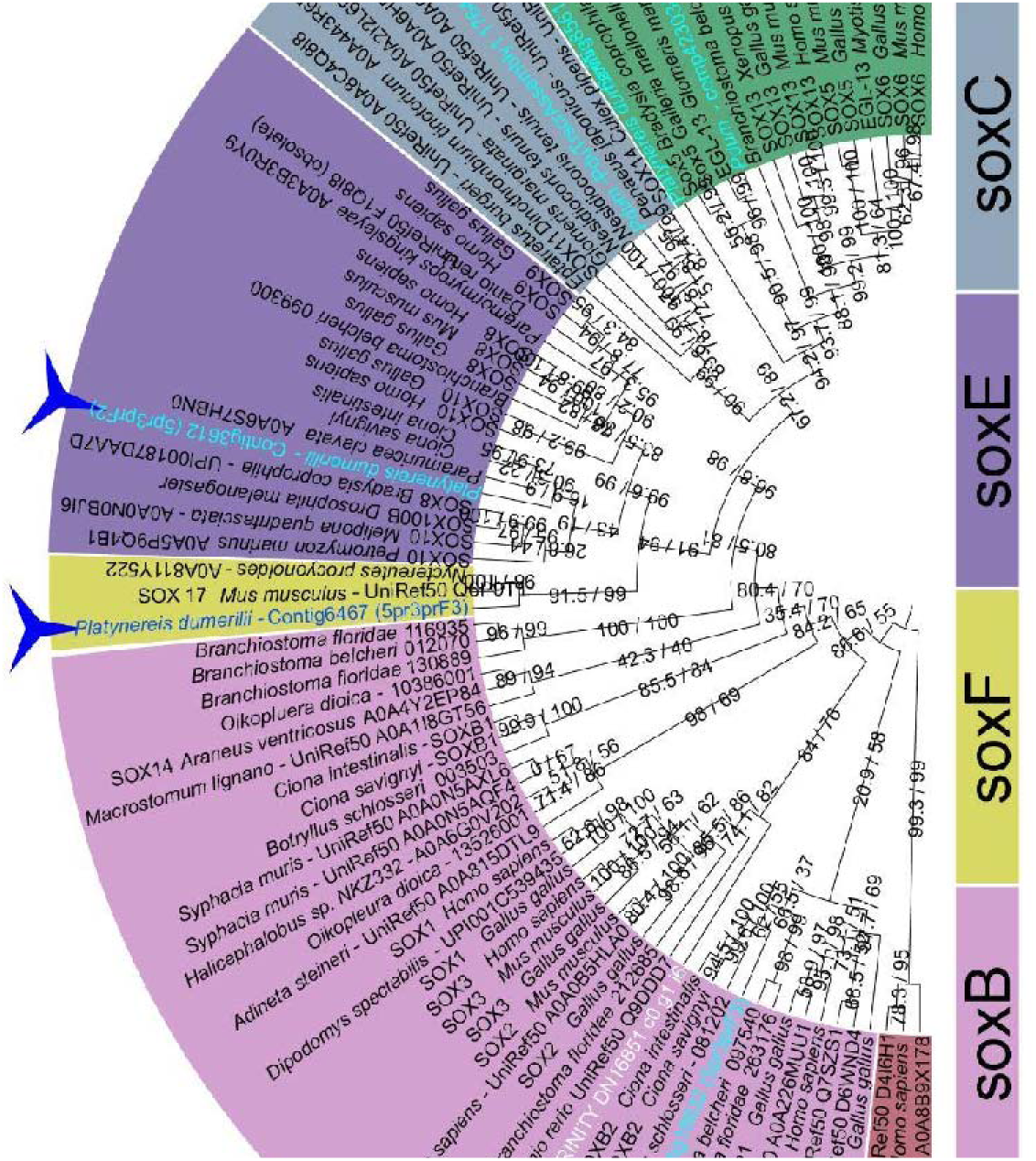
*P. dumerilii* SOX gene tree. Star highlights *sox2* transcript found to be slightly up-regulated in females according to our RNAseq analysis. Approximate maximum likelihood gene tree of SOX family members including *Platynereis* orthologs obtained from IQTree2. Protein homology was ascribed based upon their co-clustering human, *Drosophila*, and mouse proteins. SOX2 is a member of the larger SOXB family. Blue shapes (three- and five-pointed stars) at the perimeter indicate *Platynereis dumerilii* sequences. The five-pointed star at the perimeter indicates the *Platynereis dumerilii* SOX2 sequence of interest. Node support is expressed through SH-alrt and UltraFastBootstrap values as assigned by IQTree2. A high resolution version of this figure are available in our github repository (ÖzpolatLab-GitHub-Ribeiro, 2024).

**Supplementary Fig. S13.**
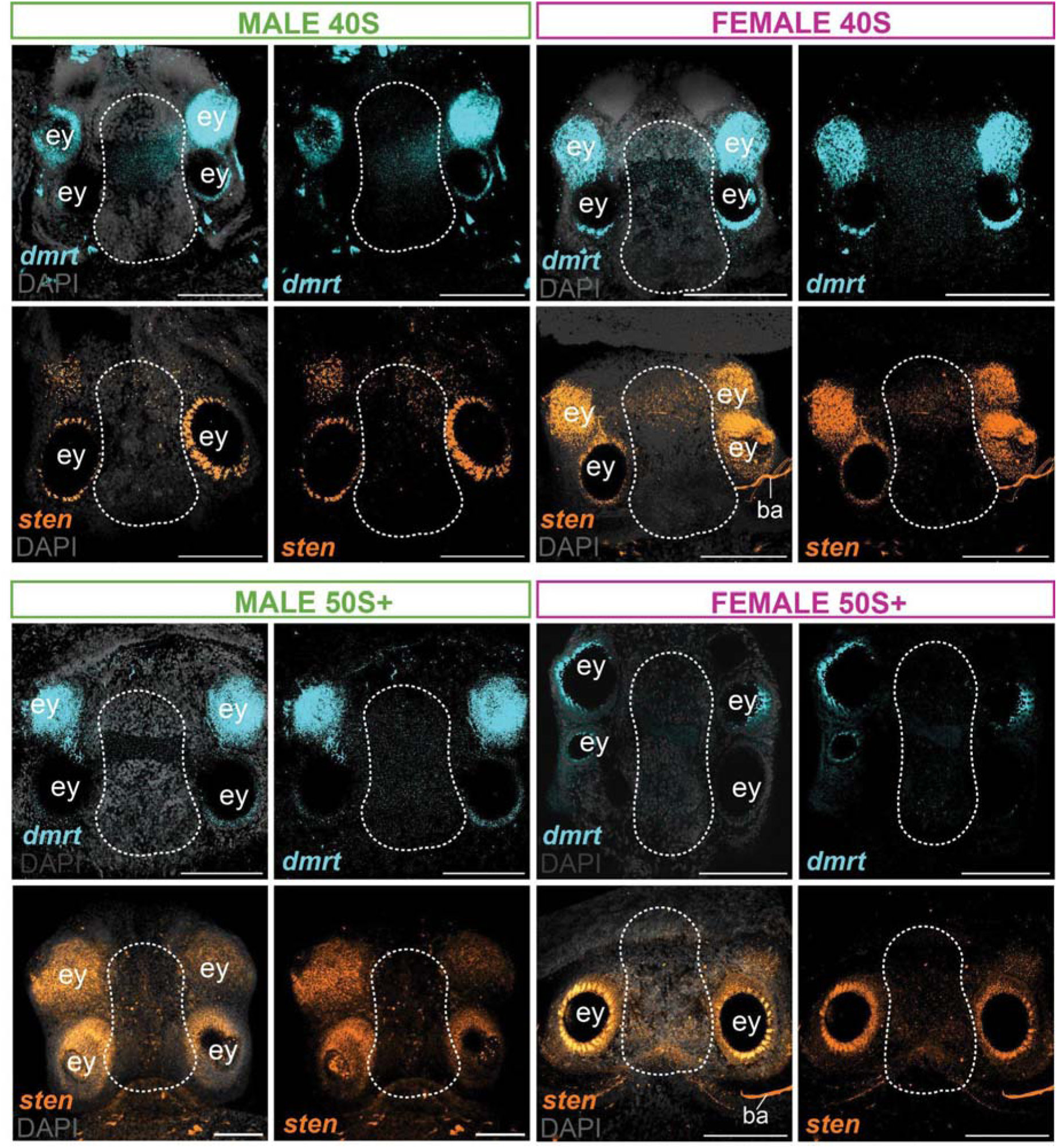
Expression of *sten* and *dmrt1* in the brains of male and female juveniles. Expression of both *dmrt1* and *sten* is generally reduced in the brain of juveniles. Dashed circles indicate the cerebral ganglion (brain). Non-specific fluorescence sometimes appears on the dorsal focal plane around the eye (ey) region in both *sten* and *dmrt1* images. Abbreviations: ey - eye. Scale bars: 100 µm.

**Supplementary Fig. S14.**
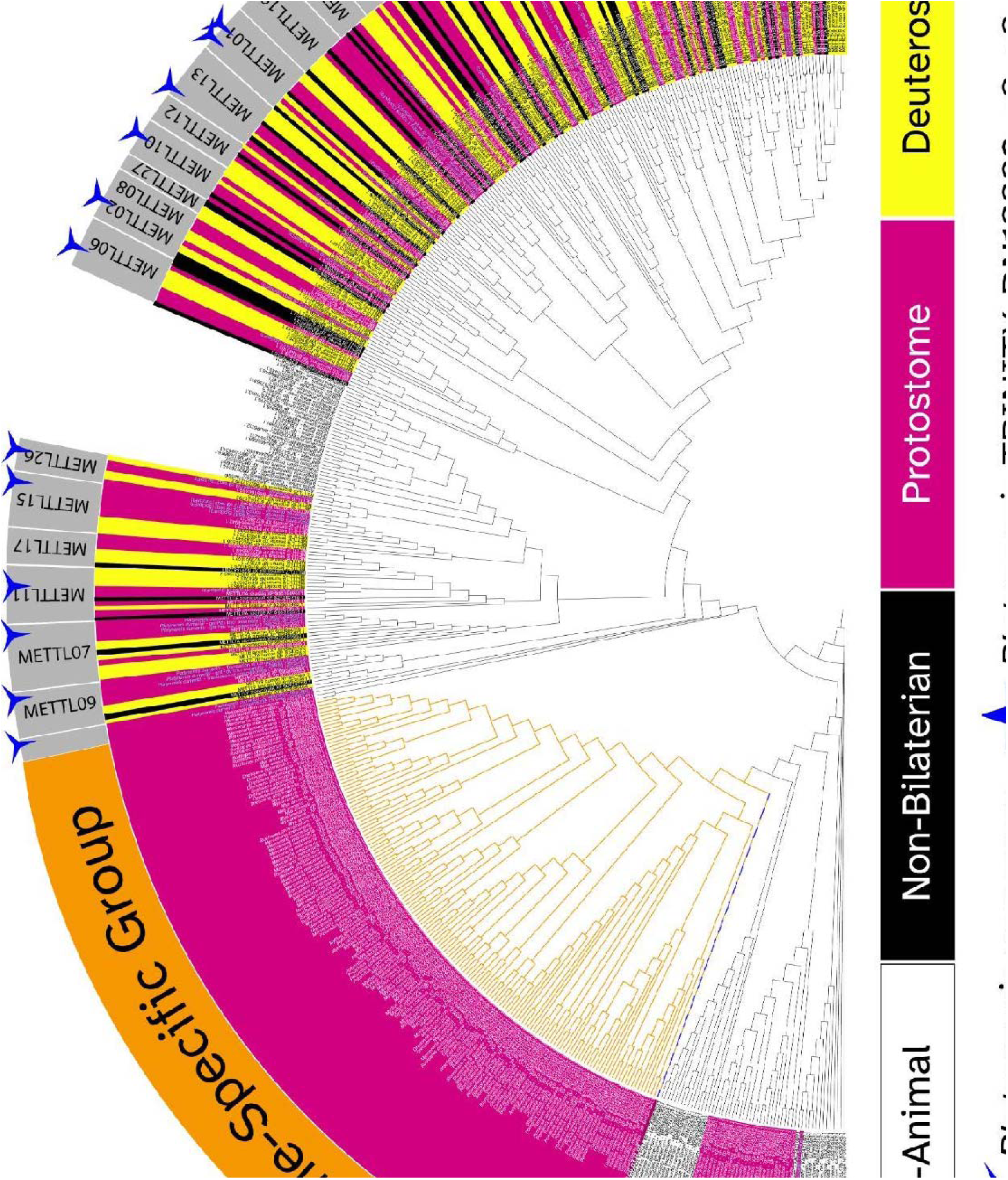
Gene tree placement of novel, protostome-specific methyltransferase (PSMT). Approximate maximum likelihood tree of Methyltransferase proteins obtained from IQTree2. Protein homology was ascribed based upon their co-clustering human, *Drosophila*, and mouse proteins (bold face IDs). Blue shapes (three- and five-pointed stars) at the perimeter indicate *Platynereis dumerilii* sequences. The 5-pointed star shows the location of the PSMT sequence identified as the differentially expressed gene of interest. Orange-stroked portion of the tree further analyzed in Supplementary Figs. S15 and S16. A gene’s background color (magenta, yellow, white, and black) indicates the taxonomic grouping of the organism from which the gene derives and is explained as follows – white: non-animal taxa, black: non-bilaterian animals, pink: Protostome bilaterians, and yellow: Deuterostome bilaterians. Protostome group contains sequences of the top hits obtained from BLAST annotation strategies, which are enriched for mollusc sequences. A high-resolution version of this tree can be found in our github repository (ÖzpolatLab-GitHub-Ribeiro, 2024).

**Supplementary Fig. S15.**
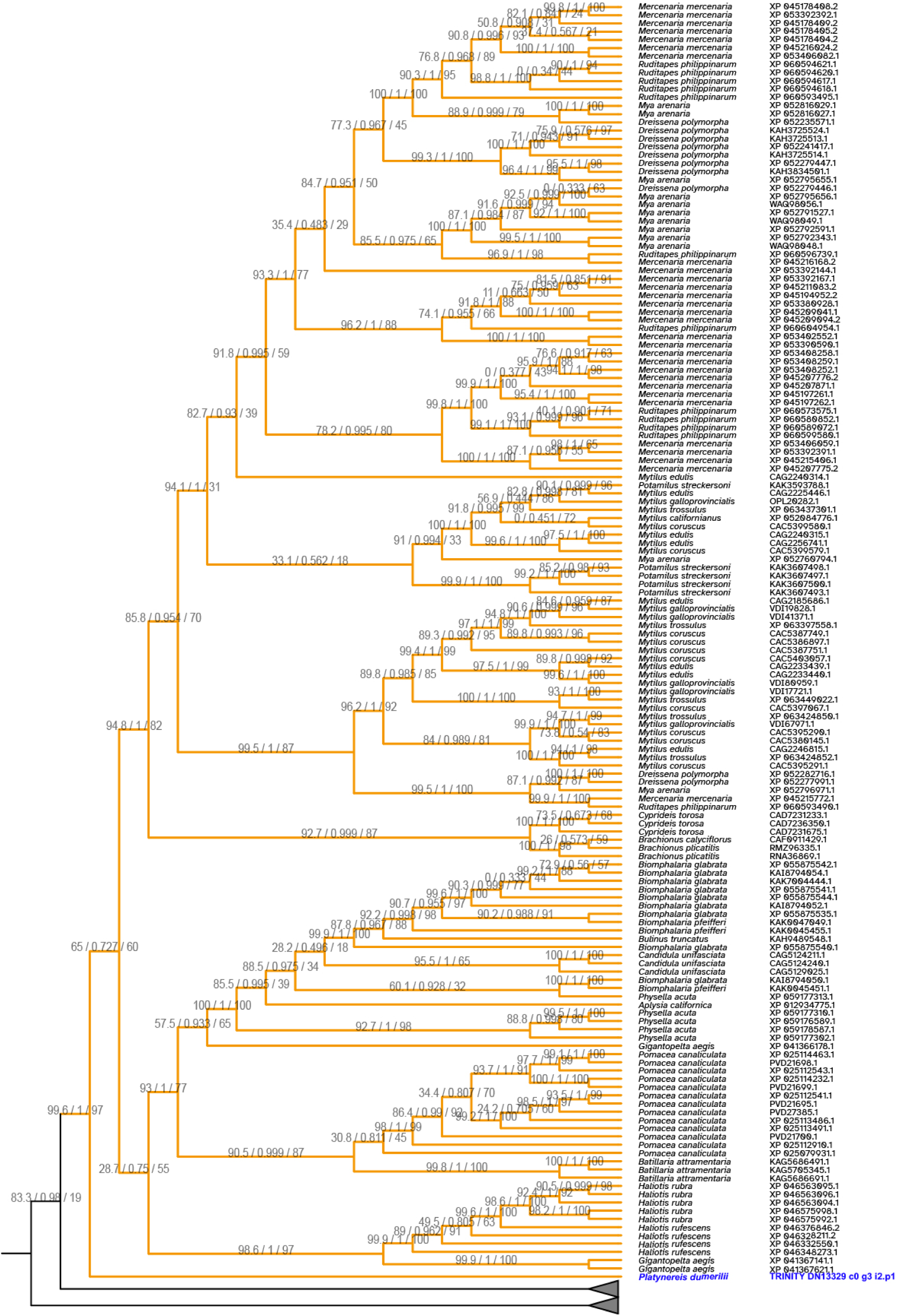
Protostome-specific methyltransferase (PSMT) Family subtree. Subtree extracted from Supplementary Fig. S13 containing the *P. dumerilii* PSMT. Support values are SH-alrt / Approximate Bayesian / UltraFastBootstrap as assigned by IQTree2. Organism and protein accession number shown for each branch in right two columns.

**Supplementary Fig. S16.**
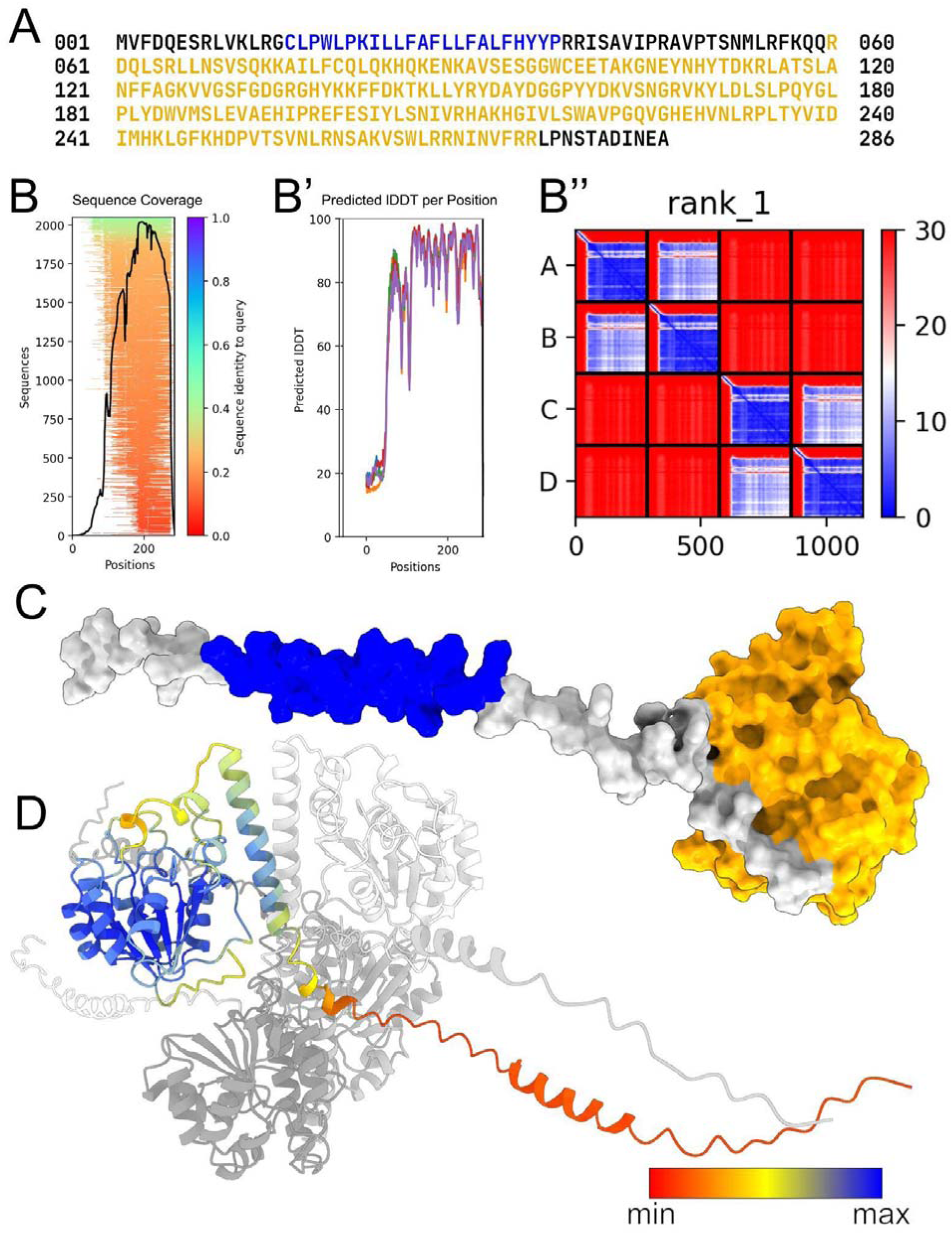
Prediction and structure comparison of novel, protostome methyltransferase. (A) Color-coded sequence of *Platynereis dumerilii* putative methyltransferase (PSMT) indicating predicted transmembrane domain (blue) and methyltransferase domain (gold). (B-B’’) ColabFold v1.5.5 reported statistics for PSMT (B) A visualization of a multiple sequence alignment shown as a line graph. The x-axis represents the 286 amino acids of the *Platynereis* PSMT sequence while the y-axis indicates the number of reference structures (y-axis) that map to the *Platynereis* residues. This graph indicates that the C-terminal region (the methyltransferase domain) of the *Platynereis* PSMT sequence had many more structures upon which its 3D structure could be modeled. (B’) Graph of Predicted local Distance Difference Test (plDDT) result following the AlphaFold prediction of the *Platynereis* PSMT. plDDT is a measure of the conservation of atomic positions between AlphaFold’s PSMT prediction and the ground truth crystal structures used to create it. plDDT is reported in the graph’s y-axis per residue along PSMT’s 286AA-long primary sequence (x-axis) with higher values suggesting more similarity between the two. This graph indicates that there is higher structural conservation in PSMT’s C-terminal methyltransferase domain (B’’) - Heatmap of the predicted aligned error (PAE). PAE is another metric of confidence for the predicted PSMT structure that compares the difference in the distance of any residue, “O”, in the model to any other residue, “J”, in the model with the distance of the same two residues (O*,J*) in the prediction. A lower PAE indicates that the distances OJ & O*J* are similar. When investigated across all possible combinations of OJ and O*J*, a 2-D matrix of values emerges, with regions of small values indicating commonly held spatial arrangements of amino acids between the reference structure and AlphaFold’s PSMT predicted structure. In this particular graph there are are multiple areas because the prediction was made as a homotetramer. Each 1/16th square of the area is a comparison of one monomer to another. As this value is a relation of position, monomer 1 will have the strongest positional correlation with monomer 1, and less with any other. Within a square, the c-terminal methyltransferase domain is most similar, even across monomers (Guo et al., 2022; Mariani et al., 2013). (C) A space-filling rendering of a monomer from the highest ranking prediction colored to indicate the transmembrane (blue) and methyltransferase domains (gold). (D) Rendering of the predicted tetrameric structure as a cartoon. One monomer has been colored based on its plDDT to show the model’s confidence and the monomer’s orientation with respect to the tetramer.

**Supplementary Fig. S17.**
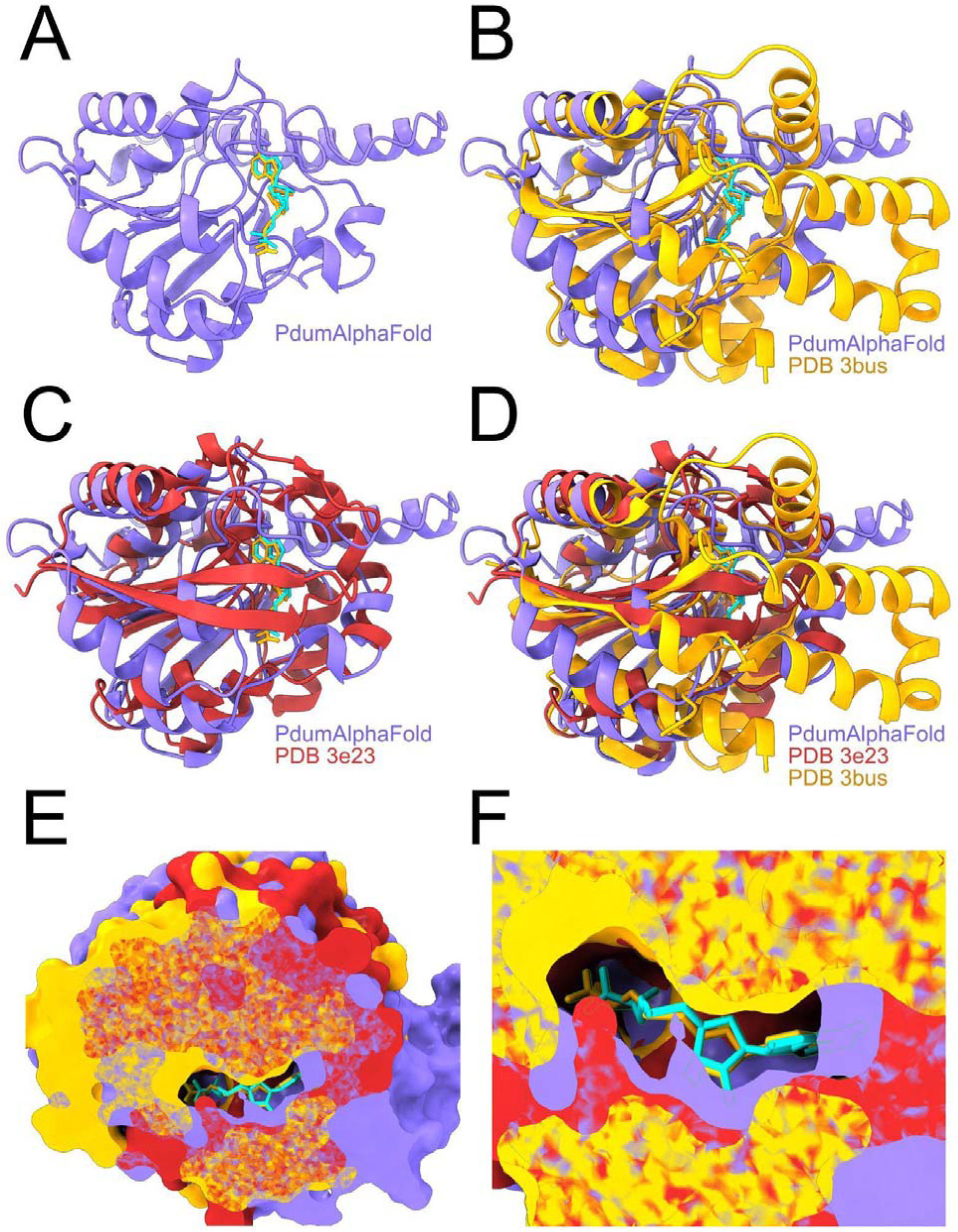
Prediction and structure comparison of PSMT. (A) ChimeraX cartoon rendering of part of the *P. dumerilii* PSMT. Predicted PSMT structure obtained using AlphaFold2 via ColabFold v1.5.5. The ligands shown are S-Adenosyl-L-Methionine (SAM) in turquoise and S-Adenosyl-Homocysteine (SAH) in yellow from Protein Data Bank (PDB) entries 3E23 and 3BUS, respectively. 3E23 and 3BUS are bona fide methyltransferase structures used by AlphaFold/ColabFold as ground-truth reference structures [refs Forouhar McCoy]. (**B-D)** ChimeraX cartoon renders of published methyltransferase crystal structures 3BUS (yellow) and 3E23 (red) overlaid upon the PSMT (purple) demonstrating similarity of the structures. (**E, F)** Space-filling render of overlaid 3E23, 3BUS, and PSMT with a cutaway section uncovering the SAM/SAH binding site, suggesting that the predicted PSMT is a bona fide methyltransferase.

## References

Altschul, S. F., Gish, W., Miller, W., Myers, E. W. and Lipman, D. J. (1990). Basic local alignment search tool. J. Mol. Biol. 215, 403–410.

Álvarez-Campos, P., Kenny, N. J., Verdes, A., Fernández, R., Novo, M., Giribet, G. and Riesgo, A. (2019). Delegating sex: differential gene expression in stolonizing syllids uncovers the hormonal control of reproduction. Genome Biol. Evol. 11, 295–318.

Balavoine, G. (2014). Segment formation in annelids: patterns, processes and evolution. Int. J. Dev. Biol. 58, 469–483.

Bar, I., Cummins, S. and Elizur, A. (2016). Transcriptome analysis reveals differentially expressed genes associated with germ cell and gonad development in the Southern bluefin tuna (*Thunnus maccoyii*). BMC Genomics 17, 1–22.

Benvenuto, C. and Weeks, S. (2020). Hermaphroditism and Gonochorism. Reprod. Biol.

Bolger, A. M., Lohse, M. and Usadel, B. (2014). Trimmomatic: a flexible trimmer for Illumina sequence data. Bioinforma. Oxf. Engl. 30, 2114–2120.

Bray, N. L., Pimentel, H., Melsted, P. and Pachter, L. (2016). Near-optimal probabilistic RNA-seq quantification. Nat. Biotechnol. 34, 525–527.

Brown, C. J., Ballabio, A., Rupert, J. L., Lafreniere, R. G., Grompe, M., Tonlorenzi, R. and Willard, H. F. (1991). A gene from the region of the human X inactivation centre is expressed exclusively from the inactive X chromosome. Nature 349, 38–44.

Bruce, H. S., Jerz, G., Kelly, S. R., McCarthy, J., Pomerantz, A., Senevirathne, G., Sherrard, A., Sun, D. A., Wolff, C. and Patel, N. H. (2021). Hybridization Chain Reaction (HCR) In Situ Protocol V.1.

Bryant, D. M., Johnson, K., DiTommaso, T., Tickle, T., Couger, M. B., Payzin-Dogru, D., Lee, T. J., Leigh, N. D., Kuo, T. H., Davis, F. G., et al. (2017). A Tissue-Mapped Axolotl De Novo Transcriptome Enables Identification of Limb Regeneration Factors. Cell Rep. 18, 762–776.

Buchfink, B., Reuter, K. and Drost, H.-G. (2021). Sensitive protein alignments at tree-of-life scale using DIAMOND. Nat. Methods 18, 366–368.

Camacho, C., Coulouris, G., Avagyan, V., Ma, N., Papadopoulos, J., Bealer, K. and Madden, T. L. (2009). BLAST+: architecture and applications. BMC Bioinformatics 10, 421.

Cantalapiedra, C. P., Hernández-Plaza, A., Letunic, I., Bork, P. and Huerta-Cepas, J. (2021). eggNOG-mapper v2: functional annotation, orthology assignments, and domain prediction at the metagenomic scale. Mol. Biol. Evol. 38, 5825–5829.

Cao, H., Li, L., Li, Z., Gao, H., Peng, G., Zhu, C., Chen, Y., Yang, F. and Dong, W. (2023). Denovo RNA-Seq analysis of ovary and testis reveals potential differentially expressed transcripts associated with gonadal unsynchronization development in *Onychostoma macrolepis*. Gene Expr. Patterns 47, 119303.

Castro-Arnau, J., Chauvigné, F., Gómez-Garrido, J., Esteve-Codina, A., Dabad, M., Alioto, T., Finn, R. N. and Cerdà, J. (2022). Developmental RNA-Seq transcriptomics of haploid germ cells and spermatozoa uncovers novel pathways associated with teleost spermiogenesis. Sci. Rep. 12, 14162.

Chadwick, D. J. and Goode, J. A. (2002). The genetics and biology of sex determination. John Wiley & Sons.

Chung, J.-J., Miki, K., Kim, D., Shim, S.-H., Shi, H. F., Hwang, J. Y., Cai, X., Iseri, Y., Zhuang, X. and Clapham, D. E. (2017). CatSperζ regulates the structural continuity of sperm Ca2+ signaling domains and is required for normal fertility. Elife 6, e23082.

Clark, W. C. (1978). Hermaphroditism as a reproductive strategy for metazoans; some correlated benefits. N. Z. J. Zool. 5, 769–780.

Clough, E., Jimenez, E., Kim, Y.-A., Whitworth, C., Neville, M. C., Hempel, L. U., Pavlou, H. J., Chen, Z.-X., Sturgill, D. and Dale, R. K. (2014). Sex-and tissue-specific functions of *Drosophila* doublesex transcription factor target genes. Dev. Cell 31, 761–773.

Crews, D. (2003). Sex determination: where environment and genetics meet. Evol. Dev. 5, 50–55.

Dong, Y., Lyu, L., Zhang, D., Li, J., Wen, H. and Shi, B. (2021). Integrated lncRNA and mRNA transcriptome analyses in the ovary of *Cynoglossus semilaevis* reveal genes and pathways potentially involved in reproduction. Front. Genet. 12, 671729.

Ebert, M. S. and Sharp, P. A. (2010). Emerging roles for natural microRNA sponges. Curr. Biol. 20, R858–R861.

Eddy, S. R. (2011). Accelerated profile HMM searches. PLoS Comput. Biol. 7, e1002195.

Engreitz, J. M., Pandya-Jones, A., McDonel, P., Shishkin, A., Sirokman, K., Surka, C., Kadri, S., Xing, J., Goren, A. and Lander, E. S. (2013). The Xist lncRNA exploits three-dimensional genome architecture to spread across the X chromosome. Science 341, 1237973.

Ewels, P., Magnusson, M., Lundin, S. and Käller, M. (2016). MultiQC: summarize analysis results for multiple tools and samples in a single report. Bioinformatics 32, 3047–3048.

Feng, B., Li, S., Wang, Q., Tang, L., Huang, F., Zhang, Z., Mahboobe, S. and Shao, C. (2021). lncRNA DMRT2-AS acts as a transcriptional regulator of dmrt2 involving in sex differentiation in the Chinese tongue sole (*Cynoglossus semilaevis*). Comp. Biochem. Physiol. B Biochem. Mol. Biol. 253, 110542.

Fischer, A. (1974). Stages and stage distribution in early oogenesis in the annelid, *Platynereis dumerilii*. Cell Tissue Res. 156, 35–45.

Fischer, A. (1975). The structure of symplasmic early oocytes and their enveloping sheath cells in the polychaete, *Platynereis dumerilii*. Cell Tissue Res. 160, 327– 343.

Fischer, A. and Hoeger, U. (1993). Metabolic links between somatic sexual maturation and oogenesis in nereid annelids—a brief review. Invertebr. Reprod. Dev. 23, 131–138.

Fischer, A. H., Henrich, T. and Arendt, D. (2010). The normal development of *Platynereis dumerilii* (Nereididae, Annelida). Front. Zool. 7, 31–31.

Gardiner-Garden, M., Ballesteros, M., Gordon, M. and Tam, P. P. (1998). Histone- and protamine-DNA association: conservation of different patterns within the β- globin domain in human sperm. Mol. Cell. Biol. 18, 3350–3356.

Gaudet, P., Škunca, N., Hu, J. C. and Dessimoz, C. (2017). Primer on the gene ontology. Gene Ontol. Handb. 25–37.

Gayen, S., Maclary, E., Hinten, M. and Kalantry, S. (2016). Sex-specific silencing of X-linked genes by Xist RNA. Proc. Natl. Acad. Sci. 113, E309–E318.

Gazave, E., Béhague, J., Laplane, L., Guillou, A., Préau, L., Demilly, A., Balavoine, G. and Vervoort, M. (2013). Posterior elongation in the annelid *Platynereis dumerilii* involves stem cells molecularly related to primordial germ cells. Dev. Biol. 382, 246–267.

Goddard, T. D., Huang, C. C., Meng, E. C., Pettersen, E. F., Couch, G. S., Morris, J. H. and Ferrin, T. E. (2018). UCSF ChimeraX: Meeting modern challenges in visualization and analysis. Protein Sci. 27, 14–25.

Gouveia-Oliveira, R., Sackett, P. W. and Pedersen, A. G. (2007). MaxAlign: maximizing usable data in an alignment. BMC Bioinformatics 8,.

Grabherr, M. G., Haas, B. J., Yassour, M., Levin, J. Z., Thompson, D. A., Amit, I., Adiconis, X., Fan, L., Raychowdhury, R., Zeng, Q., et al. (2011). Full-length transcriptome assembly from RNA-Seq data without a reference genome. Nat. Biotechnol. 29, 644–652.

Grabherr, M. G., Haas, B. J., Yassour, M., Levin, J. Z., Hompson, D. A., Amit, I., Adiconis, X., Fan, L., Raychowdhury, R., Zeng, Q., et al. (2013). Trinity: reconstructing a full-length transcriptome without a genome from RNA-Seq data. Nat. Biotechnol. 29, 644–652.

Graf, M., Teo Qi-Wen, E.-R., Sarusie, M. V., Rajaei, F. and Winkler, C. (2015). Dmrt5 controls corticotrope and gonadotrope differentiation in the zebrafish pituitary. Mol. Endocrinol. 29, 187–199.

Gu, L., Liu, H., Gu, X., Boots, C., Moley, K. H. and Wang, Q. (2015). Metabolic control of oocyte development: linking maternal nutrition and reproductive outcomes. Cell. Mol. Life Sci. 72, 251–271.

Guo, H.-B., Perminov, A., Bekele, S., Kedziora, G., Farajollahi, S., Varaljay, V., Hinkle, K., Molinero, V., Meister, K. and Hung, C. (2022). AlphaFold2 models indicate that protein sequence determines both structure and dynamics. Sci. Rep. 12, 10696.

Haas, B. J., Papanicolaou, A., Yassour, M., Grabherr, M., Blood, P. D., Bowden, J., Couger, M. B., Eccles, D., Li, B., Lieber, M., et al. (2013). De novo transcript sequence reconstruction from RNA-seq using the Trinity platform for reference generation and analysis. Nat. Protoc. 8, 1494–1512.

Hallgren, J., Tsirigos, K. D., Pedersen, M. D., Almagro Armenteros, J. J., Marcatili, P., Nielsen, H., Krogh, A. and Winther, O. (2022). DeepTMHMM predicts alpha and beta transmembrane proteins using deep neural networks. BioRxiv 2022–04.

Hart, M. W. and Foster, A. (2013). Highly expressed genes in gonads of the bat star Patiria miniata: gene ontology, expression differences, and gamete recognition loci. Invertebr. Biol. 132, 241–250.

Heenan, P., Zondag, L. and Wilson, M. J. (2016). Evolution of the Sox gene family within the chordate phylum. Gene 575, 385–392.

Hildreth, P. E. (1965). Doublesex, a recessive gene that transforms both males and females of *Drosophila* into intersexes. Genetics 51, 659.

Hu, P., Liu, B., Ma, Q., Liu, S., Liu, X. and Zhuang, Z. (2019). Expression profiles of sex-related genes in gonads of genetic male *Takifugu rubripes* after 17β-estradiol immersion. J. Oceanol. Limnol. 37, 1113–1124.

Jha, A. N., Hutchinson, T. H., Mackay, J. M., Elliott, B. M., Pascoe, P. L. and Dixon, D. R. (1995). The chromosomes of *Platynereis dumerilii* (Polychaeta: Nereidae). J. Mar. Biol. Assoc. U. K. 75, 551–562.

Ji, Y., Shen, X., Tian, S., Liu, H., Cao, T., Wang, Q. and Wang, Y. (2023). Identification, evolution and expression analysis of dmrt genes in polychaetes. Russ. J. Genet. 59, 1–8.

Jiang, Y., Sun, X., Huang, Z., Li, Z., Xu, X., Wang, W., Sun, G., Li, Y., Li, B. and Feng, Y. (2024). Transcriptome sequencing reveals the characteristics of spermatogenesis and testis development in *Amphioctopus fangsiao*. Aquac. Rep. 35, 101957.

Juliano, C. E., Swartz, S. Z. and Wessel, G. M. (2010). A conserved germline multipotency program. Development 137, 4113–4126.

Jumper, J., Evans, R., and, et al. (2021). Highly accurate protein structure prediction with AlphaFold. Nature 596, 583–589.

Jurkowski, T. P. and Jeltsch, A. (2011). On the evolutionary origin of eukaryotic DNA methyltransferases and Dnmt2. PLoS ONE 6,.

Katoh, K., Rozewicki, J. and Yamada, K. D. (2019). MAFFT online service: multiple sequence alignment, interactive sequence choice and visualization. Brief. Bioinform. 20, 1160–1166.

Kijima, T., Kurokawa, D., Sasakura, Y., Ogasawara, M., Aratake, S., Yoshida, K. and Yoshida, M. (2023). CatSper mediates not only chemotactic behavior but also the motility of ascidian sperm. Front. Cell Dev. Biol. 11,.

Kikuchi, M. and Tanaka, M. (2022). Functional modules in gametogenesis. Front. Cell Dev. Biol. 10, 914570.

Kobayashi, Y., Nagahama, Y. and Nakamura, M. (2012). Diversity and plasticity of sex determination and differentiation in fishes. Sex. Dev. 7, 115–125.

Kuehn, E., Stockinger, A. W., Girard, J., Raible, F. and Özpolat, B. D. (2019). A scalable culturing system for the marine annelid *Platynereis dumerilii*. PLoS One 14, e0226156.

Kuehn, E., Clausen, D. S., Null, R. W., Metzger, B. M., Willis, A. D. and Özpolat, B. D. (2022). Segment number threshold determines juvenile onset of germline cluster expansion in *Platynereis dumerilii*. J. Exp. Zoolog. B Mol. Dev. Evol. 338, 225–240.

Langmead, B., Trapnell, C., Pop, M. and Salzberg, S. L. (2009). Ultrafast and memory-efficient alignment of short DNA sequences to the human genome. Genome Biol. 10, R25–R25.

Leonard, J. L. ed. (2018). Transitions Between Sexual Systems: Understanding the Mechanisms of, and Pathways Between, Dioecy, Hermaphroditism and Other Sexual Systems. Cham: Springer International Publishing.

Li, W. and Godzik, A. (2006). Cd-hit: a fast program for clustering and comparing large sets of protein or nucleotide sequences. Bioinformatics 22, 1658–1659.

Li, J., Ma, W., Zeng, P., Wang, J., Geng, B., Yang, J. and Cui, Q. (2015). LncTar: a tool for predicting the RNA targets of long noncoding RNAs. Brief. Bioinform. 16, 806–812.

Lindeman, R. E., Murphy, M. W., Agrimson, K. S., Gewiss, R. L., Bardwell, V. J., Gearhart, M. D. and Zarkower, D. (2021). The conserved sex regulator DMRT1 recruits SOX9 in sexual cell fate reprogramming. Nucleic Acids Res. 49, 6144–6164.

Love, M. I., Huber, W. and Anders, S. (2014). Moderated estimation of fold change and dispersion for RNA-seq data with DESeq2. Genome Biol. 15, 1–21.

Lücht, J. and Pfannenstiel, H.-D. (1989). Spermatogenesis in *Platynereis massiliensis* (Polychaeta: Nereidae). Helgoländer Meeresunters. 43, 19–28.

Mariani, V., Biasini, M., Barbato, A. and Schwede, T. (2013). lDDT: a local superposition-free score for comparing protein structures and models using distance difference tests. Bioinformatics 29, 2722–2728.

Martin, J. L. and McMillan, F. M. (2002). SAM (dependent) I AM: the S-adenosylmethionine-dependent methyltransferase fold. Curr. Opin. Struct. Biol. 12, 783–793.

Mattick, J. S., Amaral, P. P., Carninci, P., Carpenter, S., Chang, H. Y., Chen, L.-L., Chen, R., Dean, C., Dinger, M. E. and Fitzgerald, K. A. (2023). Long non-coding RNAs: definitions, functions, challenges and recommendations. Nat. Rev. Mol. Cell Biol. 24, 430–447.

Meisel, J. (1990). Zur Hormonabhängigkeit der Spermatogenese bei *Platynereis dumerilii*: Licht- und elektronenmikroskopische Befunde sowie experimentelle Untersuchungen in vivo und in vitro.

Meng, E. C., Goddard, T. D., Pettersen, E. F., Couch, G. S., Pearson, Z. J., Morris, J. H. and Ferrin, T. E. (2023). UCSF ChimeraXL: Tools for structure building and analysis. Protein Sci. 32, e4792.

Metzger, B. and Özpolat, B. D. (2024). Developmental stage dependent effects of posterior and germline regeneration on sexual maturation in *Platynereis dumerilii*. Dev. Biol.

Minh, B. Q., Schmidt, H. A., Chernomor, O., Schrempf, D., Woodhams, M. D., von Haeseler, A. and Lanfear, R. (2020). IQ-TREE 2: new models and efficient methods for phylogenetic inference in the genomic era. Mol. Biol. Evol. 37, 1530– 1534.

Mirdita, M., Schütze, K., Moriwaki, Y., Heo, L., Ovchinnikov, S. and Steinegger, M. (2022). ColabFold: Making protein folding accessible to all. Nat. Methods 19, 679–682.

Mysore, K., Hapairai, L. K., Li, P., Roethele, J. B., Sun, L., Igiede, J., Misenti, J. K. and Duman-Scheel, M. (2021). A functional requirement for sex-determination M/m locus region lncRNA genes in *Aedes aegypti* female larvae. Sci. Rep. 11, 10657.

Nagahama, Y., Chakraborty, T., Paul-Prasanth, B., Ohta, K. and Nakamura, M. (2021). Sex determination, gonadal sex differentiation, and plasticity in vertebrate species. Physiol. Rev. 101, 1237–1308.

Nakama, A. B., Chou, H.-C. and Schneider, S. Q. (2017). The asymmetric cell division machinery in the spiral-cleaving egg and embryo of the marine annelid *Platynereis dumerilii*. BMC Dev. Biol. 17, 16.

Nawrocki, E. P. and Eddy, S. R. (2013). Infernal 1.1: 100-fold faster RNA homology searches. Bioinformatics 29, 2933–2935.

Nguyen, L.-T., Schmidt, H. A., von Haeseler, A. and Minh, B. Q. (2015). IQ-TREE: a fast and effective stochastic algorithm for estimating maximum-likelihood phylogenies. Mol. Biol. Evol. 32, 268–274.

Nieuwkoop, P. D. and Sutasurya, L. A. (1979). Primordial germ cells in the chordates: embryogenesis and phylogenesis. CUP Archive.

OpenAI (2023). ChatGPT.

Özpolat, B. D. and Bely, A. E. (2016). Developmental and molecular biology of annelid regeneration: a comparative review of recent studies. Curr. Opin. Genet. Dev. 40, 144–153.

Özpolat, B. D., Handberg-Thorsager, M., Vervoort, M. and Balavoine, G. (2017). Cell lineage and cell cycling analyses of the 4d micromere using live imaging in the marine annelid *Platynereis dumerilii*. eLife 6, e30463.

ÖzpolatLab-GitHub-Ribeiro (2024). Ribeiro-et-al_Sex_Differentiation. https://github.com/BDuyguOzpolat/Ribeiro-et-al_Sex_Differentiation.

Padilla-Gamiño, J. L., Alma, L., Spencer, L. H., Venkataraman, Y. R. and Wessler, L. (2022). Ocean acidification does not overlook sex: review of understudied effects and implications of low pH on marine invertebrate sexual reproduction. Front. Mar. Sci. 9,.

Patel, S. S., Zunjarrao, S. and Pillai, B. (2020). Neev, a novel long non-coding RNA, is expressed in chaetoblasts during regeneration of *Eisenia fetida*. J. Exp. Biol. 223, jeb216754.

Peng, J., Wei, P., Zhang, B., Zhao, Y., Zeng, D., Chen, X., Li, M. and Chen, X. (2015). Gonadal transcriptomic analysis and differentially expressed genes in the testis and ovary of the Pacific white shrimp (Litopenaeus vannamei). BMC Genomics 16, 1–18.

Penny, G. D., Kay, G. F., Sheardown, S. A., Rastan, S. and Brockdorff, N. (1996). Requirement for Xist in X chromosome inactivation. Nature 379, 131–137.

Perplexity AI (2023). Perplexity AI.

Pettersen, E. F., Goddard, T. D., Huang, C. C., Meng, E. C., Couch, G. S., Croll, T. I., Morris, J. H. and Ferrin, T. E. (2021). UCSF ChimeraXL: Structure visualization for researchers, educators, and developers. Protein Sci. 30, 70–82.

Picard, M. A.-L., Cosseau, C., Mouahid, G., Duval, D., Grunau, C., Toulza, E., Allienne, J.-F. and Boissier, J. (2015). The roles of Dmrt (Double sex/Male- abnormal-3 Related Transcription factor) genes in sex determination and differentiation mechanisms: Ubiquity and diversity across the animal kingdom. C. R. Biol. 338, 451–462.

Picard, M. A. L., Vicoso, B., Bertrand, S. and Escriva, H. (2021). Diversity of modes of reproduction and sex determination systems in invertebrates, and the putative contribution of genetic conflict. Genes 12, 1136.

Piferrer, F. (2021). Epigenetic mechanisms in sex determination and in the evolutionary transitions between sexual systems. Philos. Trans. R. Soc. B Biol. Sci. 376, 20200110.

Planques, A., Malem, J., Parapar, J., Vervoort, M. and Gazave, E. (2019). Morphological, cellular and molecular characterization of posterior regeneration in the marine annelid *Platynereis dumerilii*. Dev. Biol. 445, 189–210.

Ponz-Segrelles, G., Ribeiro, R. P., Bleidorn, C. and Molina, M. T. A. (2020). Sex-specific gene expression differences in reproducing *Syllis prolifera* and *Nudisyllis pulligera* (Annelida, Syllidae). Mar. Genomics 54, 100772.

Poon, J., Wessel, G. M. and Yajima, M. (2016). An unregulated regulator: Vasa expression in the development of somatic cells and in tumorigenesis. Dev. Biol. 415, 24–32.

Powers, N. R., Parvanov, E. D., Baker, C. L., Walker, M., Petkov, P. M. and Paigen, K. (2016). The meiotic recombination activator PRDM9 trimethylates both H3K36 and H3K4 at recombination hotspots in vivo. PLoS Genet. 12, e1006146.

Rebscher, N., Zelada-González, F., Banisch, T. U., Raible, F. and Arendt, D. (2007). Vasa unveils a common origin of germ cells and of somatic stem cells from the posterior growth zone in the polychaete *Platynereis dumerilii*. Dev. Biol. 306, 599–611.

Rebscher, N., Lidke, A. K. and Ackermann, C. F. (2012). Hidden in the crowd: Primordial germ cells and somatic stem cells in the mesodermal posterior growth zone of the polychaete *Platynereis dumerilii* are two distinct cell populations. EvoDevo 3, 9.

Ren, D., Navarro, B., Perez, G., Jackson, A. C., Hsu, S., Shi, Q., Tilly, J. L. and Clapham, D. E. (2001). A sperm ion channel required for sperm motility and male fertility. Nature 413, 603–609.

Ribeiro, R. P., Null, R. W. and Özpolat, B. D. (2024). Sexual differentiation in Platynereis dumerilii: imaging dataset. 10.5281/zenodo.11619572

Schenk, S. and Hoeger, U. (2020). Annelid coelomic fluid proteins. In Vertebrate and Invertebrate Respiratory Proteins, Lipoproteins and other Body Fluid Proteins (ed. Hoeger, U.) and Harris, J. R.), pp. 1–34. Cham: Springer International Publishing.

Schenk, S., Bannister, S. C., Sedlazeck, F. J., Anrather, D., Minh, B. Q., Bileck, A., Hartl, M., von Haeseler, A., Gerner, C. and Raible, F. (2019). Combined transcriptome and proteome profiling reveals specific molecular brain signatures for sex, maturation and circalunar clock phase. Elife 8, e41556.

Schindelin, J., Arganda-Carreras, I., Frise, E., Kaynig, V., Longair, M., Pietzsch, T., Preibisch, S., Rueden, C., Saalfeld, S., Schmid, B., et al. (2012). Fiji: An open-source platform for biological-image analysis. Nat. Methods 9, 676–682.

Schulz, G., Ulbrich, K. P., Hauenschild, C. and Pfannenstiel, H. (1989). The atokous-epitokous border is determined before the onset of heteronereid development in *Platynereis dumerilii* (Annelida, Polychaeta). Dev. Biol. 198, 29– 33.

Shen, W., Le, S., Li, Y. and Hu, F. (2016). SeqKit: a cross-platform and ultrafast toolkit for FASTA/Q file manipulation. PloS One 11, e0163962.

Shirane, K., Miura, F., Ito, T. and Lorincz, M. C. (2020). NSD1-deposited H3K36me2 directs de novo methylation in the mouse male germline and counteracts Polycomb-associated silencing. Nat. Genet. 52, 1088–1098.

Siebert, S. and Juliano, C. E. (2017). Sex, polyps, and medusae: Determination and maintenance of sex in cnidarians. Mol. Reprod. Dev. 84, 105–119.

Silva, J., Mak, W., Zvetkova, I., Appanah, R., Nesterova, T. B., Webster, Z., Peters, A. H., Jenuwein, T., Otte, A. P. and Brockdorff, N. (2003). Establishment of histone h3 methylation on the inactive X chromosome requires transient recruitment of Eed-Enx1 polycomb group complexes. Dev. Cell 4, 481–495.

Simão, F. A., Waterhouse, R. M., Ioannidis, P., Kriventseva, E. V. and Zdobnov, E. M. (2015). BUSCO: assessing genome assembly and annotation completeness with single-copy orthologs. Bioinforma. Oxf. Engl. 31, 3210–3212.

Smith, C. A. (2010). Rowley review. Sex determination in birds: a review. Emu 110, 364–377.

Speer, K. F., Allen-Waller, L., Novikov, D. R. and Barott, K. L. (2021). Molecular mechanisms of sperm motility are conserved in an early-branching metazoan. Proc. Natl. Acad. Sci. 118, e2109993118.

Steinegger, M. and Söding, J. (2017). MMseqs2 enables sensitive protein sequence searching for the analysis of massive data sets. Nat. Biotechnol. 35, 1026–1028.

Sun, Q., Huang, M. and Wei, Y. (2021). Diversity of the reaction mechanisms of SAM-dependent enzymes. Acta Pharm. Sin. B 11, 632–650.

Teufel, F., Almagro Armenteros, J. J., Johansen, A. R., Gíslason, M. H., Pihl, S. I., Tsirigos, K. D., Winther, O., Brunak, S., von Heijne, G. and Nielsen, H. (2022). SignalP 6.0 predicts all five types of signal peptides using protein language models. Nat. Biotechnol. 40, 1023–1025.

Tilic, E., Herkenrath, T., Kirfel, G. and Bartolomaeus, T. (2023). The cellular 3D printer of a marine bristle worm—chaetogenesis in Platynereis dumerilii (Audouin & Milne Edwards, 1834) (Annelida). Cell Tissue Res. 391, 305–322.

Van Blerkom, J. (2011). Mitochondrial function in the human oocyte and embryo and their role in developmental competence. Mitochondrion 11, 797–813.

Varabyou, A., Salzberg, S. L. and Pertea, M. (2021). Effects of transcriptional noise on estimates of gene and transcript expression in RNA sequencing experiments. Genome Res. 31, 301–308.

Wallis, M., Waters, P. and Graves, J. (2008). Sex determination in mammals—before and after the evolution of SRY. Cell. Mol. Life Sci. 65, 3182–3195.

Wang, Z., Qiu, X., Kong, D., Zhou, X., Guo, Z., Gao, C., Ma, S., Hao, W., Jiang, Z., Liu, S., et al. (2017). Comparative RNA-Seq analysis of differentially expressed genes in the testis and ovary of Takifugu rubripes. Comp. Biochem. Physiol. Part D Genomics Proteomics 22, 50–57.

Wang, Q., Cao, T., Wang, Y., Li, X. and Wang, Y. (2023). Genome-wide identification and comparative analysis of Dmrt genes in echinoderms. Sci. Rep. 13,.

Weeks, S. C., Benvenuto, C. and Reed, S. K. (2006). When males and hermaphrodites coexist: a review of androdioecy in animals. Integr. Comp. Biol. 46, 449–464.

Wexler, J. R., Plachetzki, D. C. and Kopp, A. (2014). Pan-metazoan phylogeny of the DMRT gene family: a framework for functional studies. Dev. Genes Evol. 224, 175–181.

Wong, J. M. and Eirin-Lopez, J. M. (2021). Evolution of methyltransferase-like (METTL) proteins in Metazoa: a complex gene family involved in epitranscriptomic regulation and other epigenetic processes. Mol. Biol. Evol. 38, 5309–5327.

Wu, S., Mipam, T., Xu, C., Zhao, W., Shah, M. A., Yi, C., Luo, H., Cai, X. and Zhong, J. (2020). Testis transcriptome profiling identified genes involved in spermatogenic arrest of cattleyak. PloS One 15, e0229503.

Yan, H., Shen, X., Cui, X., Wu, Y., Wang, L., Zhang, L., Liu, Q. and Jiang, Y. (2018). Identification of genes involved in gonadal sex differentiation and the dimorphic expression pattern in Takifugu rubripes gonad at the early stage of sex differentiation. Fish Physiol. Biochem. 44, 1275–1290.

Yan, H., Liu, Q., Jiang, J., Shen, X., Zhang, L., Yuan, Z., Wu, Y. and Liu, Y. (2021). Identification of sex differentiation-related microRNA and long non-coding RNA in Takifugu rubripes gonads. Sci. Rep. 11, 7459.

Young, M. D., Wakefield, M. J., Smyth, G. K. and Oshlack, A. (2010). Gene ontology analysis for RNA-seq: accounting for selection bias. Genome Biol. 11, 1–12.

Zarkower, D. and Murphy, M. W. (2022). DMRT1: an ancient sexual regulator required for human gonadogenesis. Sex. Dev. 16, 112–125.

Zelada Gonzáles, F. (2005). Germline development in Platynereis dumerilii and its connection to embryonic patterning. *PhD Thesis Univ. Heidelb. Ger*.

Zhang, J., Yu, P., Zhou, Q., Li, X., Ding, S., Su, S., Zhang, X., Yang, X., Zhou, W. and Wan, Q. (2018). Screening and characterisation of sex differentiation-related long non-coding RNAs in Chinese soft-shell turtle (*Pelodiscus sinensis*). Sci. Rep. 8, 8630.

Zhao, J., Sun, B. K., Erwin, J. A., Song, J.-J. and Lee, J. T. (2008). Polycomb Proteins Targeted by a Short Repeat RNA to the Mouse X Chromosome. Science 322, 750–756.

Zhu, W., Pao, G. M., Satoh, A., Cummings, G., Monaghan, J. R., Harkins, T. T., Bryant, S. V., Voss, S. R., Gardiner, D. M. and Hunter, T. (2012). Activation of germline-specific genes is required for limb regeneration in the Mexican axolotl. Dev. Biol. 370, 42–51.

